# A high-throughput phenotypic screen combined with an ultra-large-scale deep learning-based virtual screening reveals novel scaffolds of antibacterial compounds

**DOI:** 10.1101/2024.09.11.612340

**Authors:** Gabriele Scalia, Steven T. Rutherford, Ziqing Lu, Kerry R. Buchholz, Nicholas Skelton, Kangway Chuang, Nathaniel Diamant, Jan-Christian Hütter, Jerome-Maxim Luescher, Anh Miu, Jeff Blaney, Leo Gendelev, Elizabeth Skippington, Greg Zynda, Nia Dickson, Michał Koziarski, Yoshua Bengio, Aviv Regev, Man-Wah Tan, Tommaso Biancalani

**Affiliations:** Genentech, Biological Research | AI Development, South San Francisco, 94080, USA; Genentech, Infectious Diseases, South San Francisco, 94080, USA; Genentech, Small Molecules Drug Discovery, South San Francisco, 94080, USA; Genentech, OMNI Bioinformatics, South San Francisco, 94080, USA; NVIDIA Corporation, Santa Clara, 95051, USA; MILA – Québec AI Institute / Université de Montréal; Genentech, Research and Early Development, South San Francisco, 94080, USA

## Abstract

The proliferation of multi-drug-resistant bacteria underscores an urgent need for novel antibiotics. Traditional discovery methods face challenges due to limited chemical diversity, high costs, and difficulties in identifying structurally novel compounds. Here, we explore the integration of small molecule high-throughput screening with a deep learning-based virtual screening approach to uncover new antibacterial compounds. Leveraging a diverse library of nearly 2 million small molecules, we conducted comprehensive phenotypic screening against a sensitized *Escherichia coli* strain that, at a low hit rate, yielded thousands of hits. We trained a deep learning model, GNEprop, to predict antibacterial activity, ensuring robustness through out-of-distribution generalization techniques. Virtual screening of over 1.4 billion compounds identified potential candidates, of which 82 exhibited antibacterial activity, illustrating a 90X improved hit rate over the high-throughput screening experiment GNEprop was trained on. Importantly, a significant portion of these newly identified compounds exhibited high dissimilarity to known antibiotics, indicating promising avenues for further exploration in antibiotic discovery.

## MAIN

The search for novel antibiotics to fight ever-evolving drug-resistant bacterial strains remains a critical imperative in contemporary drug discovery. Traditional approaches have long relied on high-throughput screening (HTS) of small molecule libraries^1,2^, natural product mining^3,4^, directed evolution of biologics (such as AMPs)^5^, and subsequent medicinal chemistry campaigns to optimize hits for enhanced potency^6^. However, these traditional strategies are impeded by several obstacles.

For small molecule drug discovery, even the most extensive experimental screens, involving the evaluation of 10^6^ – 10^7^ molecules, barely scratch the surface of the overly vast chemical space encompassing an estimated 10^60^ possible drug-like compounds^7^. Moreover, these experimental libraries entail substantial costs in their assembly and maintenance, compounded by the need to chemically synthesize each compound. As a result, many screens yield only a handful of promising candidates or, in discouraging cases, fail to identify any compounds suitable for further lead optimization. Furthermore, many libraries contain limited chemical diversity and tend to be biased toward certain parts of the chemical space, such as known antibiotics, natural products, or past drug discovery campaigns. This makes it very challenging to identify structurally novel compounds that could potentially lead to new mechanisms of action.

These limitations have contributed to a persistent ’discovery void’ and have impaired antibiotic research in recent decades^8^. Alarming statistics further underscore the urgency of this effort, with at least 1.27 million deaths attributed directly to antibiotic resistance annually^9^. This crisis is particularly dire for gram-negative bacteria, which have not seen the introduction of a new antibiotic class in clinical practice for over half a century. These pathogens pose unique challenges, with their dual-membrane envelope acting as a formidable barrier to compound access and robust efflux systems rapidly expelling toxic molecules^10,11^.

To overcome these constraints, the adoption of virtual screening on extensive commercial repositories of small molecules has emerged as a viable solution. Over the past decades, chemical libraries, such as Enamine REAL^12^, have expanded to contain billions of compounds, providing an abundance of new chemical entities that surpass the feasible scale of direct experimental screening. Virtual screening involves using computational algorithms to assess the activity of these compounds *in silico*, facilitating the selection of a subset for in-house testing. These algorithms can be based on physical chemistry principles or employ machine learning (ML) techniques, with the latter gaining attention in antibiotic discovery due to their ability to consider whole-cell activity^13^. Notably, in a study by Stokes *et al.*^14^, a graph neural network was employed to predict antibacterial activity for *E. coli*, revealing the potential of halicin—an initially diabetes-targeted compound. Using a similar strategy, Liu *et al*.^15^ identified an antibacterial compound with narrow-spectrum activity against *Acinetobacter baumannii*. Swanson *et al*.^16^ proposed a generative approach to design molecules that inhibit the growth of *Acinetobacter baumannii*, optimizing a pre-trained deep learning oracle. Recently, Wong *et al*.^17^ demonstrated the efficacy of a graph neural network model augmented with an explainability pipeline in revealing antibacterial activity against a methicillin-susceptible strain of *Staphylococcus aureus*.

Although these studies have ignited hope in using AI to discover a new antibiotic, advancements in this field remain constrained by several factors. First, existing studies have reported a limited number of antibacterial compounds, partly due to the fact that virtual screenings often yield low hit rates. Secondly, and perhaps most significantly, the challenge lies in discovering hits with high novelty, meaning that new compounds need to be substantially structurally dissimilar to known antimicrobial molecules. Novelty is crucial as the search for new antibiotics requires exploring unique molecular scaffolds and chemotypes, considering that exhaustive efforts have already been made over many years across known ones. Yet, previous studies considered training sets with small size and limited diversity, usually spanning a few thousand compounds, which hinder the model’s ability to generalize on different parts of the chemical space.

In this work, we combine small-molecule high-throughput phenotypic screening with a deep learning-based virtual screening strategy to discover novel antibacterial compounds (**Fig. 1a**). Our high-throughput screening encompasses nearly two million small molecules, drawn from diverse regions of the chemical space distinct from known antibiotics and natural products, thereby ensuring substantial chemical diversity. Each compound was screened against *E. coli* Δ*tolC* to assess growth inhibition, revealing over five thousand antibacterial compounds, with a hit rate of 0.26%. Leveraging this dataset, we trained our deep learning model, GNEprop, to predict antibacterial activity from chemical structures. GNEprop was designed for out-of-distribution generalization and validated by characterizing structurally different activity cliffs on a holdout dataset. Using GNEprop, we conducted virtual screening on over one billion synthesizable compounds, identifying 44 thousand predicted antibacterial molecules. Subsequently, we selected 345 compounds with varying degrees of structural dissimilarity from the training set for in-house testing, leading to the discovery of 82 new antibacterial compounds (**Fig. 1b**). This represents a 90-fold increase in hit enrichment compared to the initial HTS (from 0.26% to 23.8% hit rate). Notably, one-third of the newly identified antibacterial compounds exhibit high dissimilarity to the training set (Tanimoto similarity <0.4), which we corroborate by visual inspection of the compounds on a low-dimensional manifold of chemical structures (**Fig. 1c**) and, further, through comparison with the top similar compounds from the HTS (**Fig. 1d**). Additional characterization by dose-response assays to determine IC_50_ and minimal inhibitory concentration (MIC) assays confirmed the ability of the model to identify novel, gram-negative specific, antibacterial compounds. Selection for bacteria resistant to novel compounds and genome sequencing allowed for the identification of putative molecular targets, further confirming the ability of the model to identify active compounds. Our research represents a collaborative effort between Genentech, NVIDIA, and the MILA Institute. By harnessing the capabilities of high-throughput screening, advanced deep learning modeling, and scalable computational infrastructure, our objective is to tackle the longstanding challenges associated with the discovery of novel antibiotics.

**Figure 1.**
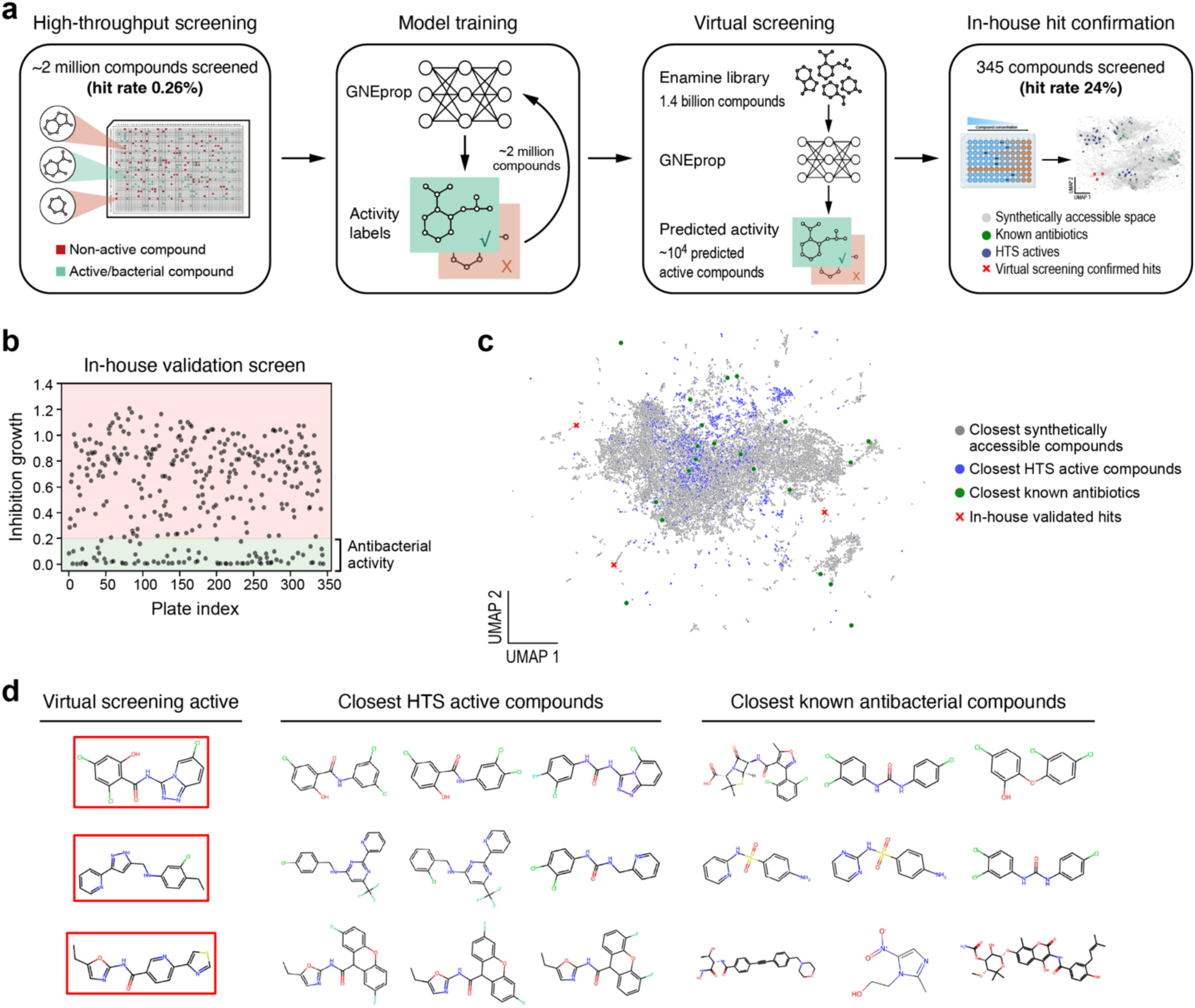
A high-throughput phenotypic screen combined with a novel deep learning strategy to enable the discovery of novel antibacterial scaffolds. **a**, we expanded the search space of novel antibacterial compounds beyond known antibiotics in two steps: a new phenotypic HTS, and a deep learning-based ultra-large-scale virtual screening of the synthesizable molecular space. The HTS enables the identification of active compounds with low similarity to known antibiotics, while covering a broad and diverse chemical space. GNEprop is a deep learning model trained on the HTS data to virtually screen the synthetically accessible space at scale. Virtual screening allows identifying novel hits with high hit rate and low similarity to HTS actives and known antibiotics, out of a large space of synthetically accessible compounds (∼10^10^). **b,** GNEprop was used to conduct a virtual screening on over 1 billion synthesizable compounds, significantly enriching for antibacterial activity (from 0.26% to 24%) compared to the HTS dataset. **c-d,** Virtual screening hits exhibit low similarity to HTS actives and known antibiotics. c, UMAP; a virtual screening hit (red cross) occupies a novel chemical space enabled by the virtual screening library, which is distinct from both HTS actives (blue dots) and known antibiotics (green dots). **d,** visual inspection of the top three Tanimoto-similar training molecules and known antibiotics for different virtual screening actives (corresponding to red crosses in panel c).

## RESULTS

### A high-throughput phenotypic screen to identify novel antibacterial compounds within families of structurally similar molecules

We initiated a phenotypic high-throughput screening (HTS) campaign using a library containing 1,981,993 small-molecule compounds (**Fig. 2a**; **Methods**). This HTS is significantly larger than datasets previously reported in the literature to train machine learning models for antibacterial property prediction^14,17–19^. Importantly, the library contains compounds that share structural similarities with known antibiotics while also exploring other regions of chemical space, thus facilitating the exploration of novel areas of the synthesizable molecular landscape (**Fig. 2a**).

**Figure 2.**
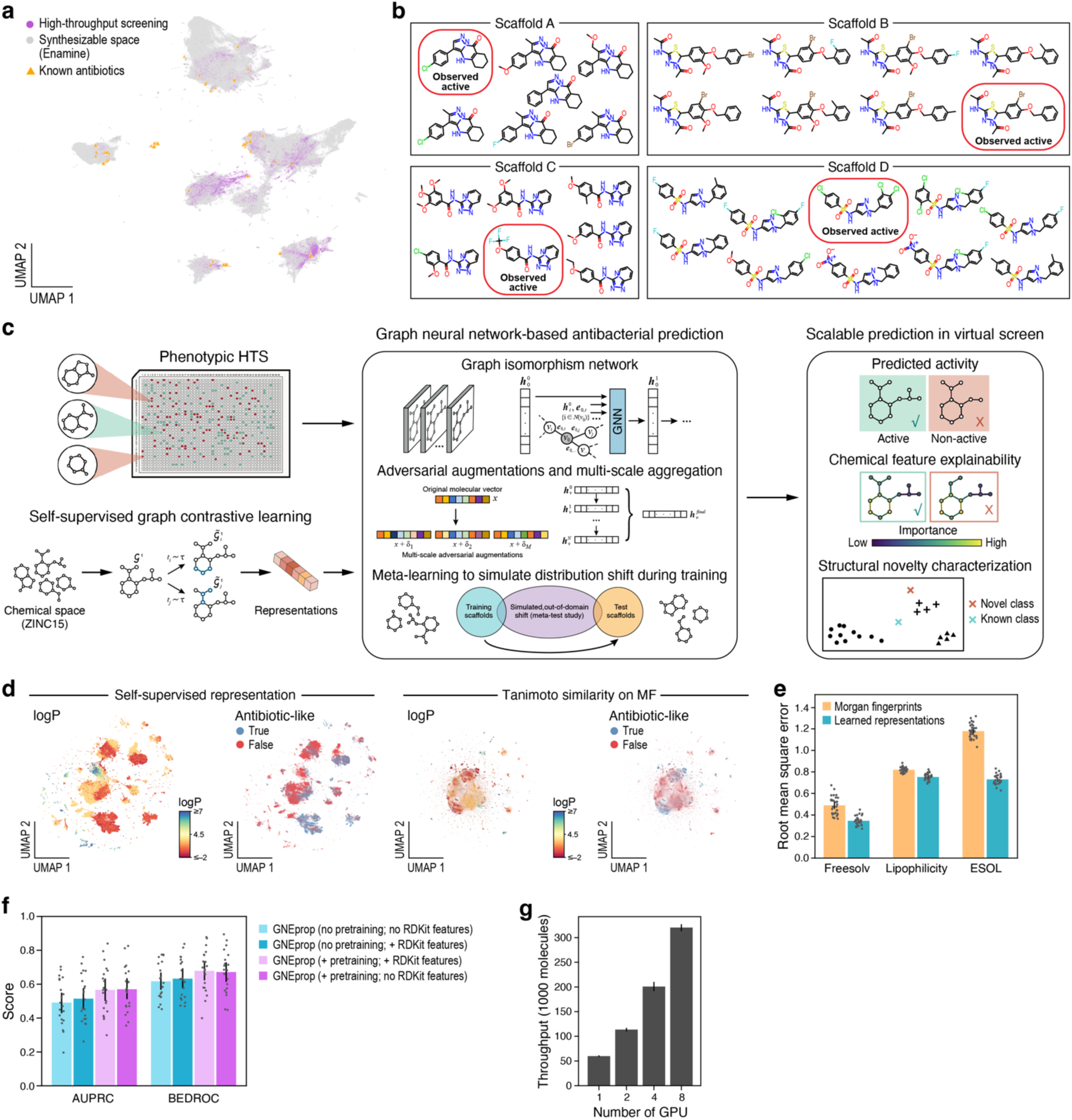
GNEprop leverages a novel HTS to explore novel parts of the chemical space at scale. **a-b**, ultra-large high-throughput phenotypic screen to identify novel and diverse antibacterial scaffolds. **a**, UMAP; the HTS (purple dots) includes compounds that have structural similarities with known antibiotics (yellow triangles), while also exploring other areas of synthesizable chemical space (grey dots), thereby enabling the exploration of new regions within the synthesizable molecular landscape. **b**, the HTS includes a large number of activity cliffs, defined as molecules sharing the same scaffold but with minute differences conferring activity. Each rectangle identifies an activity cliff. Circled compounds are active; the other compounds are inactive. **c-g**, GNEprop is a deep learning model optimized for out-of-distribution performance, efficiency, and scalability. **c**, GNEprop is a graph neural network designed to predict antibacterial activity from the graph representation of a molecule. GNEprop leverages self-supervised contrastive learning and an architecture optimized for out-of-distribution generalization. In addition to assigning an active probability between zero and one to each compound, GNEprop is equipped with an explainability pipeline to identify important parts of the molecule leading to activity and a novelty detection pipeline to characterize structural novelty. **d**, the self-supervised representation is chemically meaningful, as it exhibits well-defined, ordered regions that capture properties such as logP and antibiotic-like medicinal chemistry filter (left; additional properties shown in Extended Data Figure 2). In contrast, a baseline approach using Morgan fingerprints of compounds does not capture the same level of clustering (right). **e**, representation comparison for chemical property prediction. Root-mean-square errors over train/test folds (dots) and their average (bars) of chemical property prediction on three datasets (Freesolv, Lipophilicity, and ESOL) using a self-supervised representation (cyan) or Morgan fingerprints (yellow) as input to a linear model. **f**, GNEprop with pre-training leads to the highest performance on an antibacterial prediction task. Barplot: performance of the GNEprop model under different conditions (colors). For each condition, we ran 20 numerical experiments (dots) with an 80/20 train/test split via the scaffold-splitting approach. Bars denote standard errors. **g**, GNEprop scalability analysis. Inference throughput in terms of thousand molecules per second, varying the number of GPUs. Bars denote standard errors.

We interrogated the library with a single-point concentration (5 µM) phenotypic assay against *E. coli* Δ*tolC*, which lacks the outer membrane channel common to multiple efflux systems (**Methods**). This strain is sensitized to diverse antibiotics that are typically excluded from the wild-type *E. coli* parent^20–22^ and was chosen to increase the number of antibacterial compounds available to the model. Each compound received a binary non-/antibacterial label, based on its ability to inhibit bacterial growth (**Methods**). The combination of compound structures and labels forms our HTS dataset, which results in 5,161 antibacterial (or active) compounds (0.26% hit rate).

Next, we inspected the distribution of antibacterial compounds in the HTS data. The whole library includes numerous families of structurally similar molecules: 181,435 scaffolds are shared between more than one compound (84% of the library), and 25,437 scaffolds are shared between more than ten. Antibacterial compounds cover 3,231 scaffolds, and 1,075 scaffolds are shared by more than ten compounds. Importantly, we observed that small structural changes within the same scaffold family trigger dramatic differences in activity (examples shown in **Fig. 2b**), known as activity cliffs^23^. We hypothesized that such diversity provides the model with crucial information on how small chemical modifications affect the antibacterial properties. Moreover, we observed how potency is not trivially correlated with molecular weight and calculated lipophilicity (**Methods**), suggesting that a trained model will not bias predictions based on simple physicochemical properties. Given the size and diversity of the chemical data, we reasoned that it would allow us to develop a model that can extrapolate to new chemical spaces, including the particularly challenging task of characterizing activity cliffs within families of molecules *dissimilar* to those used to train the model.

In what follows, we present our analysis on the whole ∼2M HTS, as well as on a subset dataset of 115,519 compounds encompassing 44,852 chemical scaffolds and including 2,918 antibacterial compounds. We are releasing the model trained on both datasets; however, only the chemical structures of the subset dataset are released. To the best of our knowledge, the released dataset, which we named *GNEtolC* is one of the largest publicly released resources for antibacterial activity in small molecules, covering a diverse chemical space that includes activity cliffs and compounds dissimilar to known antibiotics. The released dataset is illustrated in **Extended Data Fig. 1** and included as **Extended Data Table 1**.

### GNEprop: a deep learning model to explore novel parts of the chemical space at scale

We used the full HTS dataset to train a novel deep learning-based model named GNEprop. GNEprop is a graph neural network trained to predict antibacterial activity from the graph representation of a molecule, assigning a probability score between zero and one to each compound (**Fig. 2c, Methods**). We designed GNEprop with the following objectives: out-of-distribution (OOD) generalization to maintain predictive accuracy in novel areas of the chemical space; sensitivity to molecular structure to discern subtle changes in molecular structures that lead to activity cliffs; scalability to handle large libraries of synthetically accessible compounds. To achieve these objectives, we leveraged recent advances in graph representation learning (**Methods**). The GNEprop encoder is based on the graph isomorphism network^24^ to derive final atom representations and leverages jumping knowledge shortcuts^25^ to create hierarchical representations of local neighborhoods. To enhance OOD generalization and robustness to activity cliffs, we augmented the training dataset through multi-scale adversarial perturbations leveraging the FLAG framework^26^. Adversarial perturbations have been shown to improve domain generalization^27^ and bypass the manual design of chemical augmentations, which remains an open challenge^28^. To further enhance model robustness to different chemical spaces, we fine-tuned the model through a meta-learning strategy that simulates domain shifts during training^29–31^. Such a strategy promotes learning features that better generalize to multiple domains (*i.e.*, different parts of the chemical spaces in our setting). Finally, we leveraged self-supervised pre-training through contrastive learning^32^ on a large set of 122 million unlabeled compounds from ZINC15^33^ to learn general, transferable representations, which provide the starting point to fine-tune the model on the labeled HTS dataset. Large-scale pre-training has been shown to improve OOD generalization in GNN-based models^24,28,34^, in addition to providing a valuable and scalable metric to navigate and represent the chemical space, as shown below.

We evaluated the model, starting from assessing the effect of pre-training on generalization and scalability. The contrastive learning strategy used to pre-train the encoder was designed to capture the similarities and differences between molecular graphs, thus learning an expressive representation without relying on manually-selected molecular properties. Indeed, we observed that the learned representation space exhibits well-defined, ordered regions, with compounds possessing similar chemical properties or satisfying the same medicinal chemistry filter grouping together (**Fig. 2d**, **Extended Data Fig. 2**). In contrast, a baseline approach using Morgan fingerprints of compounds does not capture the same level of clustering (**Fig. 2d**, **Methods**). The improved expressiveness of the self-supervised representation is further quantified in a linear evaluation setting, where a linear model is trained on top of learned representations (**Methods**). Self-supervised representations consistently outperform a Morgan fingerprint baseline, capturing physicochemical properties relevant to medicinal chemistry campaigns (**Fig. 2e, Methods**). Finally, we assessed the impact of the self-supervised representation on antibacterial activity prediction using a public dataset of 2,335 compounds that were screened for growth inhibition of *E. coli*^14^. A previous study^14^ used a hybrid deep learning model augmented with additional molecular descriptors to model inhibition growth (antibacterial compounds correspond to >80% growth inhibition). We compared augmenting with molecular descriptors to our self-supervised representation, finding that the latter significantly outperforms the first, also making molecular descriptors redundant and achieving the overall best performance (**Fig. 2f**, **Extended Data Table 2**, **Methods**). Additionally, we note that not relying on additional molecular descriptors avoids extra computation during training and inference, which is crucial to improving the scalability of GNEprop.

Next, given that screening the space of synthetically accessible compounds requires predicting billions of enumerated molecular structures^35^, we investigated model scalability. Improved scalability has been achieved through a combination of design choices, high-performance underlying libraries, and further optimizations (**Methods**). Notably, as previously discussed, GNEprop has been designed to avoid relying on ad-hoc molecular descriptors, which can be slow to compute and difficult to parallelize; instead, GNEprop amortizes the cost of computing molecular descriptors during inference for each new molecule, with the cost of pretraining GNEprop once. Additionally, optimized molecule-to-graph featurization allows fast and memory-efficient computations. Overall, GNEprop can screen thousands of molecules per second and effectively leverage a multi-GPU environment to increase throughput both for self-supervised pretraining and inference (**Fig. 2g**). For example, the model equipped with 8 A100 GPUs processes more than 300,000 molecules per second in virtual screening, which is enough to screen the billions of molecules available in existing synthetically accessible libraries in a few hours and can be further easily scaled by increasing the number of GPUs/nodes.

### GNEprop characterizes out-of-distribution activity cliffs in large phenotypic HTS data

We trained GNEprop on our HTS dataset and assessed the model’s ability to perform virtual screens by investigating prediction accuracy on out-of-distribution chemical spaces and activity cliffs on novel scaffolds. Moreover, we validated predictions by explainability on novel activity cliffs. Finally, we assessed the role of the training dataset size in enabling out-of-distribution generalization and investigated the ability of the model to efficiently drive the acquisition of new data in an active learning setting.

We start the evaluation by introducing a novel train-test split strategy. In virtual screens, it is standard to rely on scaffold splitting^36–38^, but we found this was not stringent enough, as two distinct molecular scaffolds may still be relatively similar. For this, we encoded the scaffolds using the pre-trained encoder and identified *clusters* of scaffolds by computing a *k*-nearest neighbor (*k*-NN) graph followed by clustering with a community detection algorithm^39^ (**Extended Data Fig. 3**; **Methods**). We used all compounds in one scaffold cluster as a unit for splitting the datasets, an approach we called *scaffold-cluster splitting*. Compared to other splitting strategies, scaffold-cluster splitting reduced the similarity between the test and training sets, as defined by the distribution of the maximum Tanimoto similarities of test actives to training actives, with a median similarity peak of 0.40, compared to 0.60 and 0.51 for random and scaffold splitting, respectively (**Fig. 3a**). The effectiveness of scaffold-cluster splitting is further illustrated by the substantial differences for each of a few training antibacterial compounds *vs*. each of their top five most Tanimoto-similar compounds in the test set (**Fig. 3b**). We thus used scaffold-cluster splitting as a proxy to test the ability of the model to generalize to novel chemical spaces in virtual screening. We performed an 8-fold train/validation/test split of our HTS data and trained the model (*i.e.*, fine-tuned the pre-trained model) on each fold (**Methods**, **Extended Data Table 3** includes results on the full HTS dataset, **Extended Data Table 4** includes results on the *GNEtolC* dataset). Importantly, GNEprop maintained predictive performance in the most challenging scaffold-cluster setting (AUROC 0.877 ± 0.007, AUPRC 0.126 ± 0.019), also when accounting for early recognition through the Boltzmann-enhanced discrimination of receiver operating characteristic^40^ (BEDROC 0.488 ± 0.014). To get a better understanding of the representations learned by GNEprop, we inspected the embeddings before and after fine-tuning (**Fig. 3c**): before fine-tuning, active molecules are closely embedded within their respective scaffold family (**Fig. 3c**, left) and are largely indistinguishable from negatives in the same group; by contrast, after fine-tuning, active molecules are aggregated in the same region, while preserving structural information (**Fig. 3c**; right).

**Figure 3.**
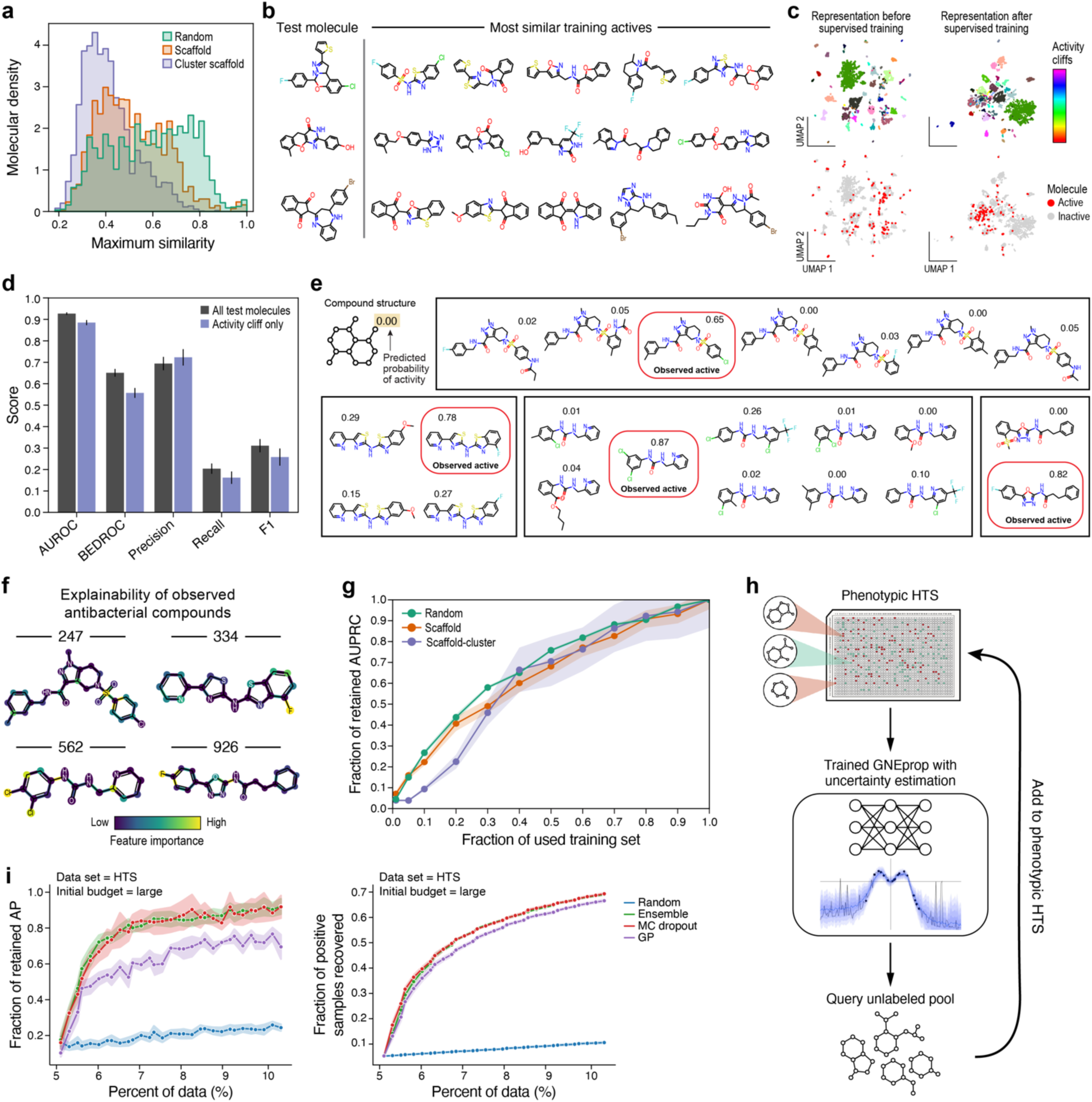
An ultra-large high-throughput phenotypic screen is combined with a deep learning model to explore novel parts of the chemical space at scale. **a**, analysis of scaffold-cluster splitting. Scaffold-cluster splitting yields train/test sets with low similarity. Probability densities for the maximum Tanimoto similarity between each test molecule vs the training set under random splitting (green), scaffold splitting (red), and scaffold-cluster splitting (purple). **b**, visual inspection of the top five Tanimoto-similar training molecules to three test molecules under scaffold-cluster splitting. **c**, visualization of activity cliffs (top) and active/inactive molecules (bottom) on the self-supervised (left) and supervised (right) representation. **d-f**, evaluation of GNEprop on activity cliffs in the HTS dataset and explainability. **d**, comparison of GNEprop performance under scaffold-cluster splitting across different metrics for all test molecules (grey bars) and activity cliff only (purple bars). Bars denote standard errors. **e**, examples of test activity cliffs correctly predicted by GNEprop. Structurally similar compounds sharing the same scaffold (i.e., activity cliffs) are grouped together (black rectangles). Active compounds are highlighted in red. The number assigned to each molecule corresponds to the predicted probability of activity. **f**, explainability of predicted active molecules for each group of the previous panel. The importance of each chemical feature (i.e., bonds and atoms) is overlayed on the compound structure (color bar). **g**, GNEprop performance (y-axis) trained on a fraction of data (x-axis) under different splitting conditions (colors). Dots denote experiments, shade area denotes standard error. **h**, active learning strategy. GNEprop extended with uncertainty estimation is used to sample new molecules to be added to the training set. **i**, the fraction of retained AP (left) and the fraction of positive samples recovered (right) as a function of the percentage of the data used in the active learning setting. Colors denote different uncertainty estimation methods, dots denote experiments and shade area denotes standard error.

We then evaluated the ability of GNEprop to predict activity cliffs, namely, chemical series sharing the same scaffolds but with minute structural changes conferring antibacterial activity. To maintain predictive power on activity cliffs, a model needs to learn small variations in input space (*i.e.*, chemical structure), which lead to large variations in target space (*i.e.*, activity). This task is particularly challenging when activity cliffs belong to families of molecules *dissimilar* to those used to train the model, as in our scaffold-cluster splitting setting. To quantitatively evaluate the performance on activity cliffs, we restricted the test set to include only structural groups with activity cliffs (**Methods**). Notably, the model’s performance is only slightly lower on the subset including activity cliffs (AUROC 0.797 ± 0.016) than that obtained on the entire test set (AUROC 0.877 ± 0.007; **Fig. 3d, Extended Data Table 5**). Further insight is provided by visualizing families of structurally similar test compounds containing an activity cliff, corroborated by their predicted probability of antibacterial activity (**Fig. 3e**).

To further quantify the relative contribution of each chemical feature (atoms and bonds) to activity, which is essential both for identifying the pharmacophore and for further medicinal chemistry optimization, we equipped GNEprop with an explainability pipeline leveraging integrated gradients^41^ (**Methods**). Explainability highlights parts of the molecule that are the most relevant for the model’s prediction of antibacterial activity. For each activity cliff correctly predicted by the model, we visualized the explanation of the antibacterial compounds as a structural heatmap, where atoms/bonds were colored by their importance for the prediction (**Fig. 3f** shows explanations for the activity cliffs in **Fig. 3e**). Explainability maps on activity cliffs can be manually validated by comparing them to the key structural features distinguishing real antibacterial compounds from the rest of the inactive compounds sharing the same scaffold in the test set. This knowledge was not available to the model, because the entire family was not used during training. Yet, we observed that in most cases, our importance maps were coherent with biochemical mechanisms that can potentially explain activity cliffs, thus supporting the model’s prediction.

To conclude the retrospective validation, we investigated the role of the HTS dataset size in out-of-distribution generalization, and how the trained GNEprop model can support the acquisition of new data. Unsurprisingly, the amount of data used for training the model has a significant impact on its final performance (**Fig. 3g**), with a larger drop in the scaffold-cluster splitting setting than under other splitting strategies for training size <40%. Crucially, this underlines the fundamental importance of the novel HTS dataset to GNEprop’s predictive performance. Notably, model performances did not plateau when the entire dataset was used, indicating that our architecture could benefit from additional data. Therefore, we investigated whether GNEprop can be used to support the acquisition of new training data. In combination with the other experiments, this analysis helps validate GNEprop predictions. Toward this goal, we evaluated GNEprop in an active learning (AL) setting (**Fig. 3h, Methods**). In this setting, GNEprop is extended with uncertainty estimation, and different techniques have been implemented to select molecules predicted to improve the model’s performance most significantly when added to the training set (**Methods**). Notably, we observed how uncertainty-guided data acquisition (in particular, based on Monte Carlo Dropout and Deep Ensembles^42^) significantly outperforms random acquisition (**Fig. 3i left**), allowing recovering 90% (compared to 20% with random sampling) of original performance with as little as 10% of the data. Importantly, we observe that AL sampling prioritizes less-represented positive samples: in earlier iterations, AL could produce over 20 times as many positive samples without any prior information about the molecule’s label (**Fig. 3i right**). Nonetheless, even in the setting where all positives are used in the initial training set, uncertainty estimation methods still outperform random sampling (**Extended Data Fig. 4**), suggesting that improvements are not purely given by higher prevalence. Overall, this analysis highlights how GNEprop could be used to drive the extension of the HTS dataset, thereby providing additional validation of its capability to retrospectively predict antibacterial activity and to improve the antibiotic drug discovery process overall.

### A novel ultra-large-scale virtual screening of antibiotics leads to improved hit rate and diversity

After evaluating the performance of GNEprop retrospectively on the HTS data, we turned to its application in a novel virtual screening setting (**Fig. 4a**).

We trained GNEprop on the entire ∼2M HTS dataset, using scaffold-cluster splitting with 95/5 for training validation. Eight replicates of the model have been trained, resampling the validation set in each, thus improving robustness and enabling uncertainty estimates^42^ (**Methods**). We used the trained model to screen the “REAL Database”, an enumerated collection of synthetically accessible compounds provided by Enamine^12^. For this study, we used a collection of approximately 1.4 billion compounds (**Methods**). As shown in **Fig. 2a**, the REAL Database provides access to a different chemical space than the HTS data and known antibiotics. Scaling GNEprop across 64 A100 GPUs (8 nodes), we were able to screen the whole space for all replicates in less than 48 hours. All compounds in the REAL Database were ranked by their predicted antibacterial activity, and those with an ensemble score greater than 0.6 were selected as hits predicted by the virtual screens (“virtual hits”), corresponding to 44,437 compounds (distribution of scores shown in **Fig. 4a, middle**). For each virtual hit, we computed its maximum structural similarity to both training set antibacterial compounds and known antibiotics, observing a distribution of maximum similarities that covers a broad range of values, with virtual hits reporting a maximum similarity as low as 0.21 (**Fig. 4b, left**), with a similarity to known antibiotics as low as 0.11 (**Fig. 4b, right**) (**Methods**).

**Figure 4.**
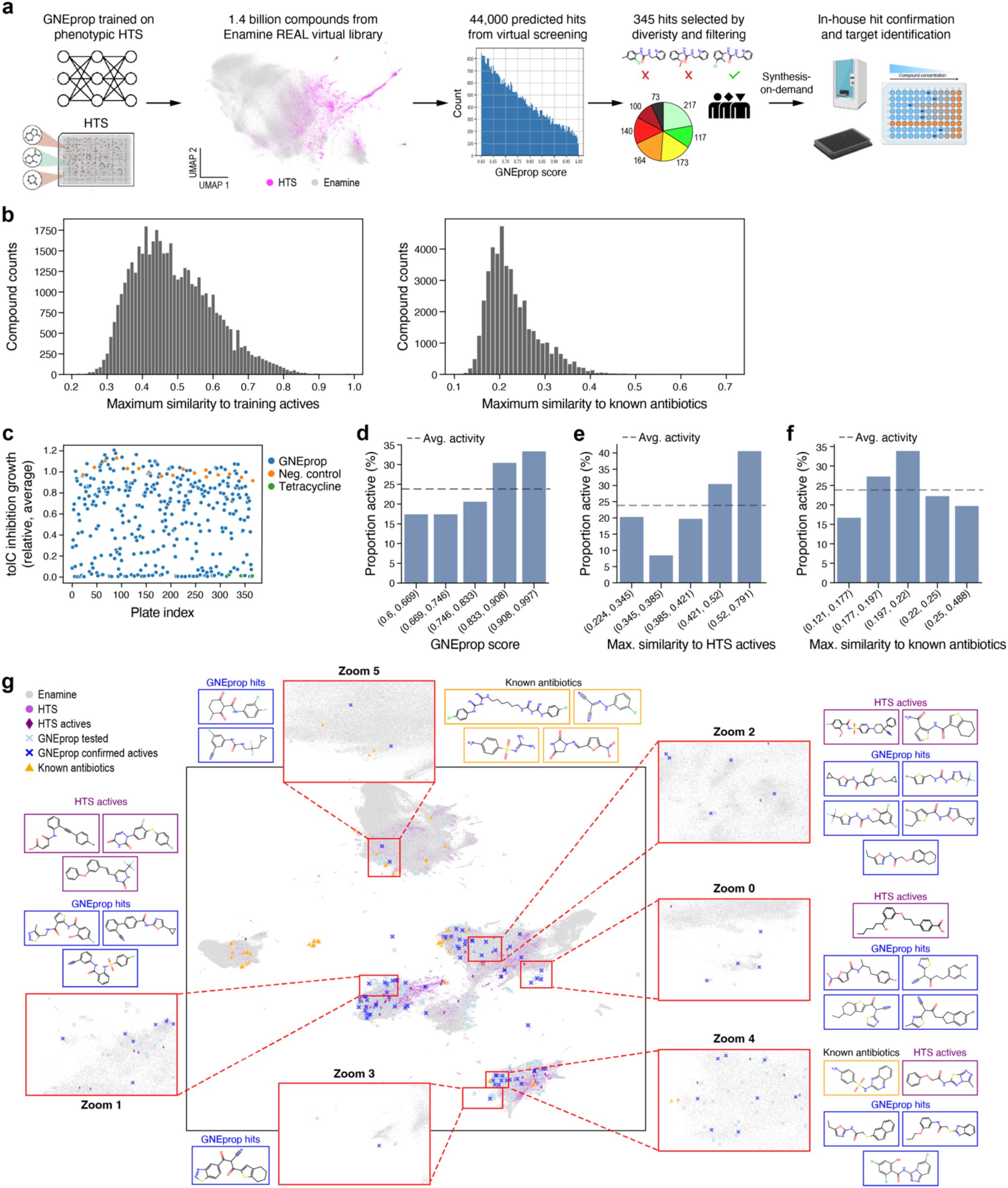
An ultra-large-scale virtual screening of antibiotics leads to novel, diverse hits. **a**, virtual screening pipeline. The GNEprop model trained on the HTS dataset is used to screen the synthetically accessible space (Enamine). Virtual hits are ranked by probability of activity and then further filtered based on multiple criteria, improving diversity and privileging virtual hits with low similarity to known antibiotics and the HTS. Finally, the selected library is synthesized for the hit confirmation and target identification phases. **b**, distribution of maximum similarity to training actives (left) and to known antibiotics (right) for virtual hits. **c**, relative inhibition growth of the selected library (blue dots), compared to tetracycline (green dots) and negative control (orange dots). **d-f**, hit rate analysis. In each plot, dashed line denotes average. **e**, hit rate as a function of activity probability as predicted by GNEprop. **e**, hit rate as a function of the maximum Tanimoto similarity to (training) HTS actives. **f**, hit rate as a function of maximum Tanimoto similarity to known antibiotics. **g**, UMAP. Visualization of the chemical space. Each dot is a molecule colored by the source (legend, **Methods**). Representations are based on self-supervised GNEprop representations and Euclidean distance. Insets are zoom-in visualizations. For each inset (red rectangle), the molecules (excluding the Enamine background) are shown; the bounding boxes of the molecules have the same color as the respective dots (legend). Known antibiotics and GNEprop hits are randomly subsampled for visualization purposes.

Virtual hits were further processed to reduce their number and increase diversity, exploring different hypotheses. In particular, we grouped them into 984 clusters, selecting the molecules with the highest GNEprop score in each cluster (**Methods**). Finally, this set was further filtered to account for the experimental budget, resulting in a library of 503 compounds. The final set was selected to prioritize structures with high GNEprop scores and varying similarity to the antibacterial compounds in the training set (49% of structures have distance ≤ 0.4 to training actives). 345 molecules were successfully synthesized and brought in-house for experimental validation.

The inhibitory activity of selected compounds against *E. coli ΔtolC* was first determined in a single-point phenotypic growth inhibition assay (see **Methods**). 82 compounds out of 345 were active (*i.e.*, they achieve at least 80% inhibition growth), which corresponds to a 23.8% hit rate (**Fig. 4c**; **Methods**). Compared to the initial HTS hit rate of 0.26%, our deep learning approach led to an ∼90-fold improvement. While acknowledging the differences in experimental design between the two campaigns, this result represents a significant advancement in the practicality and feasibility of successfully identifying initial hits. Examining the confirmed hits, we observed that GNEprop probability scores weakly correlate with the inhibition growth (Spearman correlation=0.22, P=3.4e-05), suggesting that the model has learned to predict inhibition levels, even if provided with binary labels during training. Notably, high-score predictions are significantly enriched for antibacterial activity, with predictions with a score greater than 0.9 reporting a hit rate of 36% (**Fig. 4d**). Predictive performance decreased but remained significant on molecules with low similarity to active molecules in the training set: confirmed positives are found for the lowest tested similarities (**Fig. 4e**), with 29.2% of the confirmed hits (corresponding to 24 compounds) having maximum similarity to known antibacterial compounds lower than 0.4, and the minimum similarity being 0.22. In particular, the hit rate on the subset of compounds with low maximum similarity to HTS actives (i.e., ≤ 0.4; 169 molecules in total) is 14.2%, which, albeit being lower than the overall hit rate (23.8%), is significantly higher than the initial HTS dataset (0.26%). Compared to known antibiotics, the vast majority of the confirmed positives show substantial novelty, with 98.8% of the confirmed antibacterial compounds having low maximum similarity to known antibiotics (i.e., ≤ 0.4) and 39% having very low similarity (i.e., ≤ 0.2). Interestingly, we found that the hit rate is not strongly affected by the similarity to known antibiotics (**Fig. 4f**), likely because the HTS dataset also samples from regions of the chemical space which are quite distinct from those occupied by known antibiotics. Overall, these results suggest that even though the performance of the model decreases as the distance from the training HTS increases, (1) the model retains a significantly higher hit rate even for molecules dissimilar to HTS actives, and (2) the vast majority of the confirmed hits are structurally novel compared to known antibiotics. These observations are further substantiated by a UMAP visualization (**Methods**), which illustrates that confirmed hits (blue crosses) are reasonably distinct both from the HTS (purple dots, with HTS actives as dark-purple diamonds) and known antibiotics (yellow triangles) (**Fig. 4h**; insets highlight specific examples). The full list of confirmed hits is included in **Extended Data Table 6**.

### Biological characterization of novel compounds confirms antibacterial activity

While the single-point growth inhibition assay validated the selection of antibacterial compounds using GNEprop, we carried out additional experiments to further confirm and characterize the activity of the identified compounds, investigate activity on additional bacterial strains, and identify the targeted protein or pathway.

First, we selected all compounds with >50% inhibition against *E. coli* Δ*tolC* (165 compounds), which was a larger set than the compounds that we initially defined as active (>80% inhibition). We performed a dose-response assay on these compounds to identify the half-maximal inhibitory concentration (or IC_50_), which denotes the concentration at which growth is inhibited by 50% (**Fig. 5a**). The resulting IC_50_ values ranged from <1 µM to greater than 40 µM **(Fig. 5b**), indicating 95% (157) of the compounds initially tested have a measurable IC_50_ value on *E. coli* Δ*tolC*. IC_50_ were also measured against wild-type MG1655 *E. coli*, which is efflux-proficient and has an intact outer membrane, and the gram-positive bacterium *Staphylococcus aureus*. Additionally, as an initial test to identify generally lytic molecules, the dose-response assay was run against a yeast isolate, *Saccharomyces cerevisiae* (**Methods**). This removed 6 of the 165 compounds from further investigation (**Extended Data Table 7**).

**Figure 5.**
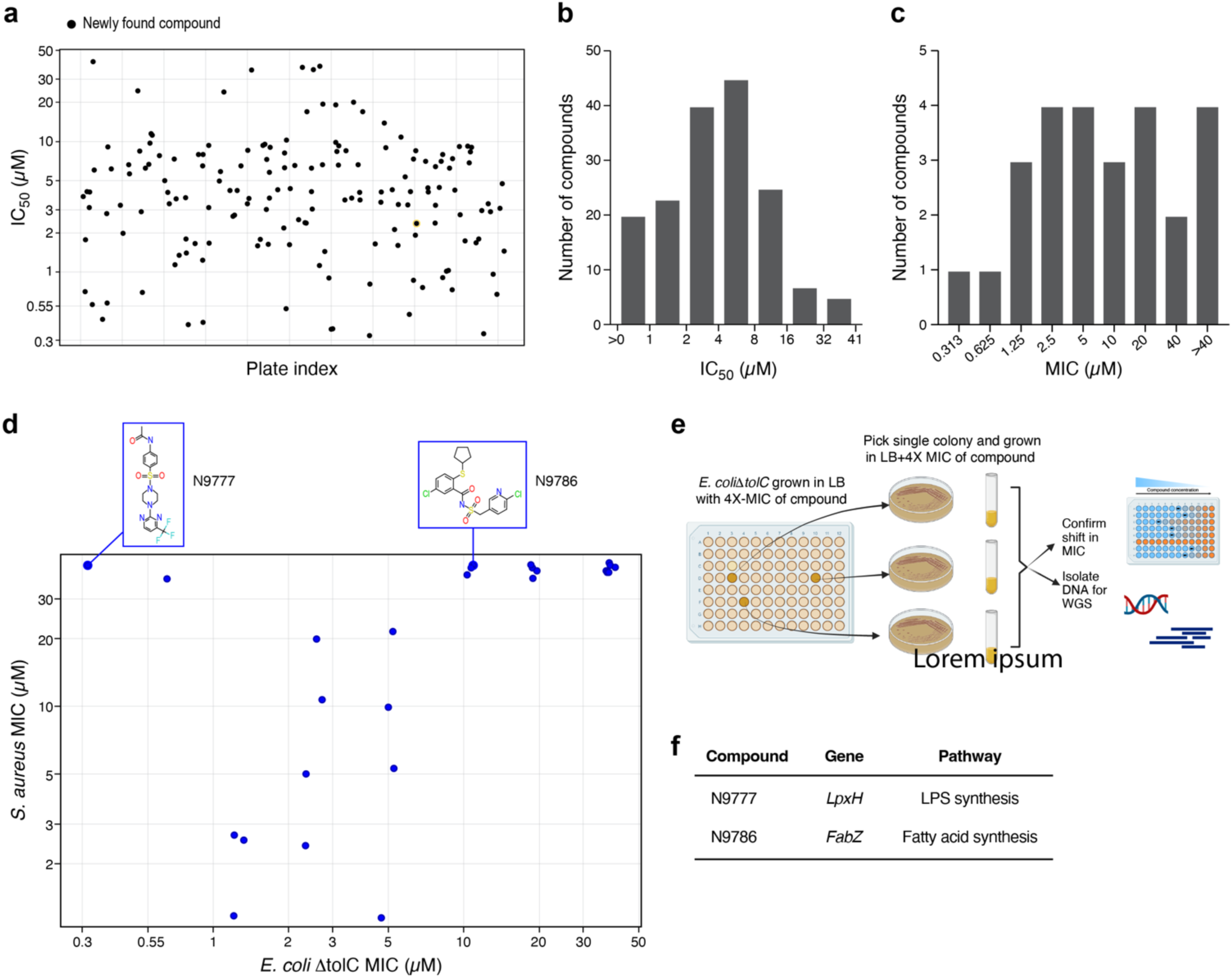
Biological characterization of virtual screening hits confirms antibacterial activity. **a**, IC_50_ values of compounds tested in dose-response assay against *E. coli* Δ*tolC*, x-axis, compound index. **b**, distribution of IC_50_ values. **c**, distribution of minimal inhibitory concentration (MIC) assay values against *E. coli* Δ*tolC*, see also Extended Data Table 6. **d**, visualization of selected molecules as a function of *E. coli* Δ*tolC* activity (MIC, x-axis) and *S. aureus* activity (MIC, y-axis). Structures of compounds selected for target identification are shown in blue rectangles. **e**, cartoon schematic of selection of resistant clones. **f**, summary of results from whole genome sequencing of resistant clones, see also Extended Data Table 8.

Next, we performed a minimal inhibitory concentration (MIC) assay to identify the lowest dose at which complete inhibition of bacterial growth was observed after overnight incubation with compound, which is a more stringent and universal test of antibiotic activity than the IC_50_. A total of 26 compounds were selected from the dose-response data using the following criteria: *E. coli* Δ*tolC* <5 µM and 3-fold selective over *S. aureus*, suggesting they are gram-negative selective; those that were most potent on *E. coli* Δ*tolC* (IC_50_ <0.03 µM); and those with inhibitory activity against wild-type *E. coli* (IC_50_ <10 µM). Twenty-two of the compounds tested against *E. coli* Δ*tolC* had MICs ranging from 0.3 to 40 µM (**Fig. 5c, Extended Data Table 8**). Interestingly, there was an overall trend for molecules with more potent activity against the *E. coli* Δ*tolC* mutant to have higher GNEprop scores (Spearman correlation between MIC and GNEprop score = 0.26), even though GNEprop was not trained on MIC or other dose-dependent assays. Thus, using GNEprop predictions, we were able to quickly identify novel compounds with whole-cell MICs.

Next, we tested the compounds against wild-type *E. coli* in the MIC assay. We reasoned that even though GNEprop was trained with data from a screen using the *E. coli* Δ*tolC* mutant, compounds with activity against wild-type *E. coli* could potentially be a subset of these hits. Here, none of the tested compounds exhibited a MIC against bacteria with an intact outer membrane, demonstrating the limitation of training with data from a sensitized *E. coli* strain for identifying molecules to further develop into antibiotics and this further highlights the barrier to gram-negative antibiotic discovery^43^. Future efforts will focus on replicating the HTS and deep learning pipeline on wild-type gram-negative strains, thus potentially enabling the discovery of candidates suitable for further development.

The compounds active against *E. coli* Δ*tolC* likely include both molecules with gram-negative selective activity as well as broad spectrum molecules with activity against gram-negative and gram-positive bacteria. To assess this, we tested the same set of 26 active compounds for MICs against *Staphylococcus aureus*, a gram-positive bacterium, and found that 58% (15 compounds) of the hits we characterized exhibited MICs against both the *E. coli* Δ*tolC* mutant and *S. aureus* (**Extended Data Table 8**). The broad-spectrum activity is not due to general lytic properties of the compounds as they did not show activity in a red blood cell hemolysis assay (**Extended Data Fig. 5**). Thus, GNEprop was able to identify both gram-negative specific and broad-spectrum antibacterial molecules.

Finally, we focused on gram-negative selective molecules to identify whether they can identify specific gram-negative targets in *E. coli*. We selected two compounds, labeled N9777 and N9786, that exhibited a gram-negative-selective spectrum with MICs of 0.313 µM and 10 µM, respectively, against the *E. coli* Δ*tolC*, and MICs >40 µM against the gram-positive *S. aureus* (**Fig. 5d** and **Extended Data Table 8**). These compounds lacked wild-type *E. coli* activity but did not lyse red blood cells, suggesting that they hit specific essential targets unique gram-negative bacteria but are unable to overcome tolC-mediated efflux.

To identify the targets, we selected spontaneous resistant mutants to N9777 and N9786 using *E. coli* Δ*tolC* (**Fig. 5e**). For N9777, we isolated two independent clones with 16 to >128-fold increases in MIC compared to the parent strain. Whole genome sequencing of these two clones revealed two distinct mutations in the *lpxH* gene, which encodes an essential enzyme in the lipopolysaccharide (LPS) biosynthesis pathway^44^ (**Fig. 5f, Extended Data Table 9**). Confirming this, a structurally similar compound (AZ^45^) has been reported to inhibit another gram-negative bacterium, *Klebsiella*, and the *lpxH* N9777-resistance mutations we isolated map to the identified AZ binding pocket on *Klebsiella* LpxH. Selections with N9786 resulted in eight resistant clones with 4 to 8-fold higher MICs compared to the *E. coli* Δ*tolC* parent strain. Whole genome sequence analysis identified six distinct mutations in *fabZ* and one mutation in *acpP* (**Fig. 5f, Extended Data Table 9**). Both genes encode enzymes with roles in fatty acid biosynthesis: FabZ is an essential dehydratase^46^, and AcpP is the carrier protein for the growing fatty acid chain^47^. While fatty acid biosynthesis is not specific to gram-negatives, it does form essential precursors for LPS synthesis, which is essential in *E. coli* and specific to gram-negative bacteria^48^. Thus, antibacterial molecules discovered with our virtual screen identify specific, validated targets in gram-negative bacteria.

### Towards improved characterization of the cell envelope barrier and novel mechanisms of action in gram-negative bacteria

Our virtual screening strategy primarily focused on the early *hit identification* phase, providing a diverse and novel set of candidate antibacterial compounds that have been further assayed to determine their mechanism of action. Our HTS dataset utilized a Δ*tolC* mutant *E. coli* strain, which is defective for efflux of toxigenic molecules, including antibiotics. As previously discussed, this choice reduced false negatives by ensuring that potential antibiotics that might otherwise be subject to efflux are included and increased the number of antibacterial compounds available to train the deep learning model. However, it also prevents screening for cell membrane permeability, which needs to be experimentally assayed. In the following, we performed additional screening against *E. coli* wild-type and used the differential labels to build a predictive model for cell membrane permeability. Finally, we extend our virtual screening strategy with an anomaly detection pipeline to aid in the early identification of novel MoA of the hits.

To investigate the ability to separately model cell membrane permeability, we screened 3,074 of the 5,161 previously identified Δ*tolC E. coli* antibacterial compounds (based on compound availability) against the wild-type *E. coli* strain. We identified 306 molecules (∼10%) active against both mutant and wild-type, whereas 2,768 were active only against the mutant. We leveraged this dataset to evaluate the ability of GNEprop to distinguish antibacterial compounds that overcome the permeability barrier. In particular, we re-trained GNEprop on the entire HTS dataset with three class labels: inactive, active against *E. coli* Δ*tolC* only (label: "inhibiting"), and active against both *E. coli* Δ*tolC* and wild-type *E. coli* (label: "cell-penetrating"). We used an eight-fold 80/10/10 train/validating/test split with stratified sampling and class-weighted cross-entropy loss to account for highly imbalanced data (**Methods**). GNEprop maintained predictive power in this setting (**Fig. 6a**). However, while the fraction of molecules misclassified inactive was similar between the two strains, it was more challenging to correctly classify cell-penetrating compounds: of the 122 compounds classified as active (inhibiting or cell-penetrating), only half were correctly classified as cell-penetrating. We hypothesized that the limited number of labeled cell-penetrating compounds and the fact that inhibiting and cell-penetrating compounds share the same chemical space in the self-supervised representation contribute to the difficulty (**Extended Data Fig. 6**).

**Figure 6.**
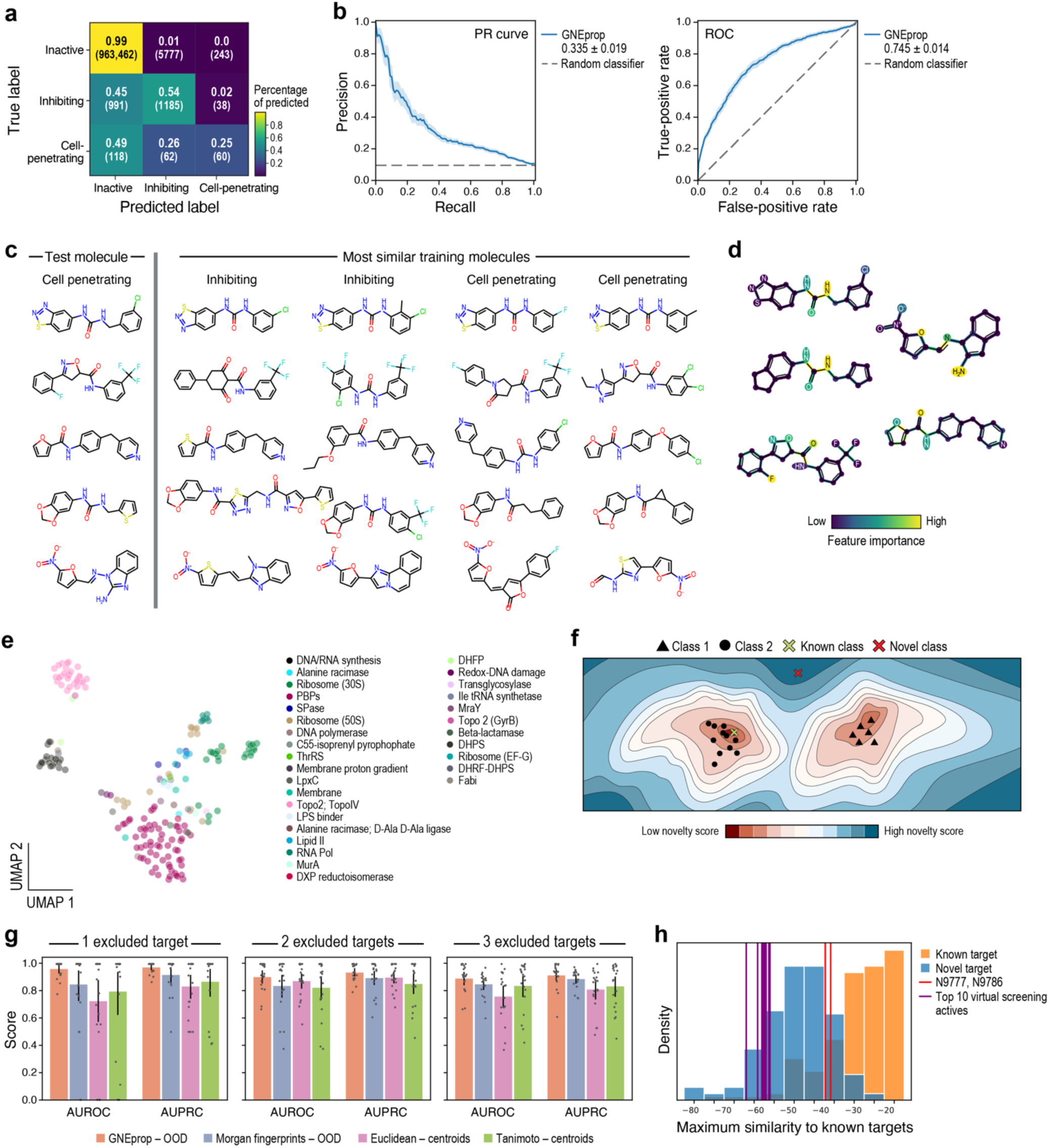
Exploring GNEprop for cell envelope barrier characterization and novel mechanisms of action prediction in gram-negative bacteria. **a-d**, GNEprop, combined with a double-assayed HTS, improves the characterization of the cell envelope barrier. **a**, GNEprop classifies novel compounds as inactives, inhibiting (i.e., active only against *E. coli ΔtolC*) and cell-penetrating (i.e., active against both *E. coli ΔtolC* and *E. coli* wild-type). Confusion matrix. **b**, GNEprop performance classifying cell-penetrating vs inhibiting compounds. Numbers in legend refer to the area under the curve. **c**, activity cliffs in permeability. For five cell-penetrating compounds (left column), the two most structurally similar inhibiting and cell-penetrating compounds are shown. **d**, the predicted chemical feature importance is overlayed to the chosen cell-penetrating compounds (left column of panel c). **e-h**, GNEprop supports the identification of potentially novel MoA. **e**, low dimensional plots showing known antibiotics, encoded on the self-supervised representation, colored by target. **f**, cartoon schematic. A molecule with novel MoA (red X), encoded in the self-supervised representation, is far from known target clusters. In contrast, a molecule with an existing MoA (green X) is close to a known cluster. **g**, retrospective evaluation of the pipeline to detect novel mechanisms of action. Antibiotics binding to a number of known targets (titles) are excluded from the training representation and tested whether detected novel by four methods (colors). GNEprop-OOD denotes the proposed pipeline, the other methods correspond to different baselines. Results are averaged over splits (dots). **h**, distribution of maximum similarity to known MoA, as computed by the proposed pipeline. Known targets and novel targets refer to excluded targets (3 excluded targets case), combined across multiple splits, and provide reference distributions. The scores of the top 10 virtual screening activities by distance to known targets are shown in purple. The scores of N9777 and N9786 are shown in red.

Next, by leveraging our explainability pipeline, we investigated the ability of GNEprop to identify chemical motifs linked to cell permeability. Because activity in the wild-type strain implicitly combines permeability with inhibition, the explainability framework would not disentangle features related to permeability when GNEprop is trained in a three-class setting. Therefore, we re-trained GNEprop to only differentiate between "inhibiting" and "cell-penetrating" labels (**Methods**). We notice that, despite the small size of this dataset, the model achieved significant predictive power (**Fig. 6b**, AUROC = 0.745 ± 0.014, AUPRC = 0.335 ± 0.019), also thanks to the pre-training strategy (**Methods**). Interestingly, 25% of correctly predicted "cell-penetrating" molecules have the highest similarity to a training "inhibiting" molecule. This demonstrates structural cliffs with respect to permeability (**Fig. 6c**), such that minimal structural changes in some molecules within a family of structurally similar active ones confer cell permeability. Our explainability framework then highlighted parts of the molecules that, based on the model, are important for cell membrane permeability (**Fig. 6d**). Though this data set lacks the depth to elucidate a set of general retention rules for compounds in gram-negative bacteria, we did observe instances consistent with previous reports. For example, a primary amine (**Fig. 6d**; right) was highlighted as important for the cell-penetrating activity, and this feature was previously found to be predictive for accumulation in gram-negative bacteria^49^. Although these explainability masks are difficult to validate, due to the smaller number of compounds to compare them against, the predictions remain largely consistent, suggesting that GNEprop retains predictive power in characterizing structural cliffs for permeability.

Finally, we investigated the ability of GNEprop to aid in the identification of potentially novel MoA. Leveraging recent advances in deep learning-based out-of-distribution detection^50,51^, we extended GNEprop with a novelty detection pipeline, to predict whether a molecule likely has a novel MoA compared to a set of known antibiotics. First, we assembled a dataset of 196 known antibiotics, grouped into 31 classes according to their targets (**Extended Data Table 10**). Because this dataset was small, we relied solely on our self-supervised representation, and leveraged the key observation that the embedded antibiotics partition into distinct clusters largely consistent with their target (**Fig. 6f, Methods**), outperforming using conventional Tanimoto similarity on Morgan fingerprints (**Methods, Extended Data Table 11**). Therefore, we hypothesized that antibacterial molecules embedded next to a known antibiotic have a higher likelihood of sharing the same target, whereas antibacterial molecules embedded far from any previously characterized compounds (*i.e.*, “outliers”) are more likely to exhibit a distinct MoA. This defines our approach for detecting potentially novel MoAs, devising a method to measure the latent distance between a novel compound and known classes of antibiotics. In particular, we fit class-conditional Gaussian distributions with respect to the representations learned by GNEprop for known targets to estimate a “novelty” score for new molecules based on the Mahalanobis distance (**Fig. 6f**, **Methods**). We benchmarked the proposed method retrospectively excluding a set of targets from the training set. As an ablation study, the proposed method was compared to three baselines, each following a similar strategy, but where one or more components (GNEprop representations, Mahalanobis distance, class-conditional distributions) are missing (**Methods**). On average, the proposed approach outperforms all the considered baselines across all the experiments and exhibits the lowest variance, thus highlighting its robustness (**Fig. 6g**; **Extended Data Table 12**). The proposed approach leads to a distribution of scores for novel targets significantly different from that of known targets (**Fig. 6h**, t-test P=1.4e-43). Finally, we applied the proposed pipeline to the set of 82 antibacterial compounds identified in our virtual screen. Here, consistent with the previous analysis, we found a significant fraction of antibacterial molecules with high novelty compared to known targets (top-10 antibacterial molecules by novelty score shown in **Fig. 6h**, purple lines). Even though comparatively less novel, reflecting their known targets, N9777 and N9786 also have a higher likelihood under the “novel” distribution. The novelty score only requires computing Mahalanobis distances on molecular embeddings, being therefore extremely efficient, with virtually no overhead on the virtual screening. Overall, the proposed approach has shown promise as an efficient and effective tool to aid the prioritization of novel compounds, complementing the virtual screening strategy.

## DISCUSSION

The pursuit of new antibiotics to combat the continuously evolving drug-resistant bacterial strains remains a crucial priority in modern drug discovery. Recent advances in the field of deep learning modeling have ignited hope in the search for new antibacterial compounds. In particular, virtual screenings offer the opportunity to search the chemical space for active small molecule compounds on scales too vast for experimentation. Still, progress has been limited by several factors. Virtual screenings often have low hit rates, and finding highly novel compounds is challenging. Novelty is essential, as new antibiotics must differ significantly from known antimicrobial molecules to overcome their limitations. Additionally, previous studies have leveraged small and limited training sets, restricting the models’ ability to learn complex relationships between chemical structures and growth inhibition, especially in the presence of activity cliffs and diverse scaffolds, limiting the ability to explore the broader chemical space effectively.

This work combines a diverse HTS library of nearly 2 million small molecules with a deep learning strategy and a virtual screening library of over 1.4 billion compounds to explore the chemical space at scale. Through this approach, we were able to significantly increase the hit rate and enhance the novelty and diversity of the identified hits. Overall, the framework described constitutes an end-to-end pipeline that combines the strengths of HTS and virtual screening, overcoming their individual limitations, to expand the search for novel antibacterial compounds.

Our experiments analyzing the performance of the model when varying the training set have highlighted the importance of a large and diverse training set to achieve strong out-of-distribution results. Notably, GNEprop’s performance did not plateau even when utilizing the entire dataset, suggesting the model could benefit from additional data. Importantly, our experiments using GNEprop to support the acquisition of new training data showed promising results, allowing recovering 90% of original performance with just 10% of the data. This suggests how an active learning strategy^52,53^ that leverages the deep learning model (and its uncertainty^42^) to “guide” the acquisition of new molecules over multiple iterations in a *lab-in-the-loop* setting could further accelerate the search for novel antibacterial compounds, reducing the number of experiments needed and the associated time. Future work should focus on how to effectively balance exploration and exploitation in this setting^54^, while also accounting for novelty and the desired activity profile.

Our experiments equipping GNEprop with an explainability pipeline have shown how this approach can help elucidate minute structural differences which lead to major changes in activity, as learned by the deep learning model. This has been validated not only through the comparison of active molecules to similar inactive ones, but also by differentiating molecules able to penetrate the cell membrane to similar active but not cell-penetrating ones, recapitulating known biochemistry^49^. While in this study we leveraged this approach to validate the predictions of our model, similar approaches have the potential to help the discovery of “general rules” governing activity across different strains (and differential activity between them), for example, through global explainability techniques^55,56^. This could lead to a better understanding of the biochemical features conferring activity, which can be used for improved library design, lead optimization, and biological understanding. A major challenge in this context is represented by spurious features conferring activity, exacerbated by the limited size and diversity of existing libraries. Recent work on causal representation learning^57,58^ and disentangled representations could set the basis for identifying causal or (conditionally) invariant features conferring activity.

A limitation of this study is represented by the targeted strain, *E. coli ΔtolC*, which lacks the outer membrane channel common to multiple efflux systems and is sensitized to diverse antibiotics that are typically excluded from the wild-type parent^20–22^. This choice allowed us to increase the initial number of active compounds available to the model while focusing on areas of the chemical space distant from known antibiotics and natural products. Indeed, out of nearly 2 million molecules screened, only 0.26% have been identified as active against *E. coli ΔtolC*, and our additional screening for wild-type activity on the subset of *ΔtolC* actives has reported ∼10% relative hit rate, which would lead to a ∼0.03% wild-type hit rate on the original screening. This active hit rate is much lower than that reported elsewhere, such as by Stokes et al.^14^, who screened with a wild-type *E. coli* strain. We attribute this to the higher diversity and lack of known antibiotics (or similar molecules) in the HTS library. However, the promising results of the deep learning model in the virtual screening setting, combined with its ability to differentiate between *E. coli ΔtolC* and wild-type activity in our analysis, suggest that the model could still learn robust representations in more challenging settings. Although antibacterial activity is a critical feature for discovering novel starting material, additional considerations, such as host cell toxicity and in vivo pharmacokinetics, are also important, and the proposed approach could be extended for these other activities given appropriate datasets. While this framework has been described in the context of *E. coli ΔtolC* activity, it can be easily extended to different strains and activities, which will be the subject of future work.

Our work primarily leveraged structural novelty as a proxy for biochemical novelty assuming that structurally different compounds have a higher likelihood of exhibiting a novel mode of action or different mechanisms. Indeed, as we showed in our analysis of antibiotic targets, the molecular representations learned by the model loosely cluster by known targets. This assumption often holds and is consistent with prior work on ML-driven antibiotic discovery^14,16^, and it has the main advantage of requiring only structural information as input to the model, therefore being extremely efficient in ultra-large-scale screening settings. However, a limitation of this approach is that the precise characterization of the mechanism of action (or its novelty) is often coarse-grained and imprecise. Future work could extend deep learning-based virtual screening with additional data modalities, for example, integrating phenotypic and molecular downstream effects^59^ with high-throughput and high-content technologies^60^ to better characterize MoAs and the broader functional impact of small molecules. Additionally, this work only focused on small molecule discovery. However, the search for new classes of antibacterial compounds could leverage emerging classes of therapeutic agents, such as peptides^13,61^ or other modalities^62^, which have recently found promising applications in this domain. Leveraging AI methods to integrate existing data across diverse modalities, encompassing both therapeutic agents and heterogeneous readouts, presents a challenging yet promising avenue for future research.

## METHODS

### Bacterial strains

Strains and plasmids used in this study can be found in **Extended Data Table 13**. *E. coli* K-12 MG1655 and its derivatives were used. The *E. coli* MG1655 Δ*tolC* mutant used in the phenotypic screen was previously constructed by introducing *tolC*::*aph* from the Keio collection by λ Red recombination^22,63,64^. The luminescence plasmid pRP1195^65^ was transformed into the *E. coli* MG1655 Δ*tolC* mutant by electroporation and maintained by selection with carbenicillin (50 µg/mL). The wild-type Gram-positive *Staphylococcus aureus* USA300 strain was also used in this study. All cultures were grown at 37°C in LB broth (Sigma) unless noted otherwise.

### High-throughput phenotypic screen

The set of compounds compiled for this project was screened at 5 µM for growth inhibition activity against *E. coli* MG1655 Δ*tolC* carrying pRP1195 plasmid (“*E. coli* Δ*tolC* lux”), which allowed direct readout of growth. Because the screen was carried out over multiple weeks, a single starter culture was generated to eliminate day-to-day inoculum variability. To generate the starter culture stocks, *E. coli* Δ*tolC* lux was streaked onto LB agar plates containing carbenicillin and grown at 37°C for 16 h. A single isolated colony was used to inoculate 10 mL of LB plus carbenicillin, which was grown with aeration (200 rpm shaker) at 37°C for 16 h. The culture was moved to ice for 10 min and then diluted to a final OD_600_ of 1 in ice-cold LB plus carbenicillin with 20% glycerol. Aliquots of the starter culture were stored at -80°C.

Fresh starter culture aliquots were removed from the freezer and thawed on ice each day screening was performed. The starter culture was diluted 1/1000 in fresh, chilled LB without carbenicillin. We previously confirmed that the pRP1195 plasmid was maintained in the absence of selection for the entire course of the experiment. 6 µL of the diluted starter culture was distributed into each well of 1536-well plates (Corning) using a Tempest liquid handler (Formulatrix). Wells were preloaded with screening compounds in DMSO using an Echo acoustic liquid handler (Bechman Coulter), resulting in a final concentration of 5 µM compound at <0.4% DMSO. DMSO alone (negative) or tetracycline (positive) were included as controls on every plate. Plates were incubated statically at 37°C in a humidified incubator for 6 hours, and then luminescence was read using an Envision plate reader (PerkinElmer).

Each compound received a binary score: growth inhibition of >80% (label: actives) and growth inhibition of <20% (label: inactive). Hits were subsequently confirmed by repeating the same luminescence growth inhibition assay using *E. coli* Δ*tolC* lux. Overall, this process resulted in 1,981,993 annotated compounds and a 0.26% hit rate (corresponding to 5,161 compounds).

We confirmed that potency is not trivially correlated with molecular weight and calculated lipophilicity (Pearson correlation between logP and activity: 0.019; Pearson correlation between molecular weight and activity: -0.008).

### Testing virtual screening hits for activity

Compounds identified using GNEprop model (described in following sections) were tested by pre-loading compounds into 384 well plates using Echo acoustic liquid handler for a final concentration of 10 µM or dose response described below. DMSO alone (negative) or tetracycline (positive) were included as controls on every plate. Bacterial stocks (prepared as before) were diluted to 0.001 OD_600_ in LB broth on ice, and 10 µl was added per well using a Multidrop Combi (Thermo Scientific). Plates were incubated statically at 37°C in a humidified incubator for 4 h, then 10 µl of BacTiter-Glo reagent (Promega) was dispensed into plates and luminescence was read after 10 min incubation at 37°C, using an Envision plate reader (PerkinElmer). Compounds that showed activity in the single point screen were followed up with dose-response assay in 384 well plate with eight 2-fold dilutions starting at 40 µM as the highest concentration, technical duplicates on each plate. This was repeated for a second biological replicate.

### Minimal inhibitory concentration (MIC)

A modified minimal inhibitory concentration (MIC) assay was performed by first inoculating 3 ml liquid cultures of LB with 1-2 colonies of bacteria from LB-agar plates. Tubes were grown shaking at 37°C until in log phase and diluted to 0.001 OD_600_ before 50 µl of bacteria was added to 50 µl of compound diluted in 96-well round bottom polystyrene plates (Corning) for a final volume of 100 µl. Plates were grown at 37°C, static with humidity, for approximately 18 h before the minimal concentration at which growth was inhibited was determined by visual inspection and measuring OD_600_.

### Red blood cell lysis assay

Compounds at 2x the indicated concentration were diluted in PBS in a 96-well round bottom polystyrene plates (Corning). Whole heparinized human blood was diluted and added to the compound in plate above such that the final blood concentration was 2%. Plates were incubated 37 degrees, static with humidity overnight (18h) then centrifuged at 600 x g for 3 min in a tabletop centrifuge, supernatant was removed to a new plate, careful to avoid the pellets, and OD_600_ read on a SpectraMax M5 plate reader (Molecular Devices). Percent lysis was calculated by subtracting the negative control (buffer alone) samples run in each plate from both test-compound and the 100% lysis sample (water), then calculating the amount of lysis as a percentage of the “100%” lysis sample.

### Selection of resistant clones and target ID

*E. coli* MG1655 Δ*tolC* was grown up in 4X the MIC of compound (as determined previously) in 100 µl LB in 96-well round bottom polystyrene plates (Corning) grown 37°C, static. At 24 and 48 h the plates were checked for growth, and subsequently streaked onto LB-agar plates and grown overnight 37°C. A single colony was picked and grown in the presence of compound. This culture was used to inoculate a MIC-assay as described above to confirm a shift in MIC as compared to the parental MG1655Δ*tolC,* and total DNA was extracted using the Blood and Tissue kit (Qiagen) for whole-genome Illumina sequencing. The Genomic DNA Screen Tape and Tapestation 4200 (Agilent Technologies) was used to determine the quality of the DNA and quantified using the Qubit dsDNA BR assay kit (Thermo Fisher Scientific). The Nextera DNA Flex kit (Illumina) was used for library preparation with an input of 100 ng of genomic DNA. Resulting libraries were multiplexed and sequenced on NovaSeq (Illumina), to generate 5 million, 75-bp paired-end reads for each sample.

### Variant Detection

Genomic variants were inferred from Illumina reads mapped to the E. coli K-12 MG1655 reference genome (GenBank accession no. NC_000913) using previously described methodology^66^.

### Yeast assay in 384-well format

To screen for non-specific lytic compounds in an HTS-amenable format *Saccharomyces cerevisiae* budding yeast, strain BY4741 was grown on YEPD agar plates overnight at 30 degrees. From the plate colonies were diluted into YEPD borth to an OD_600_ of 0.025 and grown shaking at 30 degrees approximately 6 hours until and OD_600_ of 0.3-0.4. This culture was diluted to 0.02 in YEPD and 10 µl per well added to 384 well plates made with the Echo acoustic liquid handler as described above for testing for bacterial growth inhibition. Geneticin at 200 ug/ml was used as a control for the yeast assay. Plates were incubated at 37 degrees static for 15 hours, then 10 µl of BacTiter-Glo reagent (Promega) was dispensed per well and luminescence was read after 10 min incubation at 37°C with the Envision plate reader (PerkinElmer).

### GNEprop model

#### Graph Encoder Architecture

The GNEprop encoder leverages Graph Isomorphism Network (GIN) layers^67^, with the extension to edge features^24^. Initial node features ℎ_*v*_(#) encode atom descriptors (*e.g.*, atomic number, valence, formal charge, etc.); end edge *e*_*uv*_features encode bond features (*e.g.*, bond type, ring membership, stereo configuration, etc.), with initial feature dimensions of 133 and 12 for atoms and bonds, respectively. Before the graph learning layers, atoms and bonds were independently encoded in a *d*_*node*_-dimensional representation (hyperparameter values are given in the "Dataset analysis" section). Atom representations were then created through *K* layers following the update function:

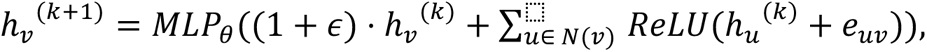

where ℎ_*v*_^(*k*)^ is a *d*_*node*_-dimensional vector denoting the representation of the *v* node in the *k* layer, *∊* is a trainable parameter, *MLP_θ_* is a 2-layer feed-forward neural network with parameter *θ* and hidden dimension 2 ⋅ *d*_*node*_, *ReLU* is the rectifier function and *N*(*v*) is the neighborhood of node *v*.

To improve the expressiveness of the embeddings and encode molecular features at different resolutions, Jumping Knowledge Networks^25^ were used. Different aggregation schemes were evaluated, with *concatenation* aggregation attaining best results. The final node representations, ℎ_*v*_*^final^*, were computed as the concatenation of [ℎ_*v*_^(1)^, …, ℎ_*v*_^(*k*)^]. Node representations were then aggregated for graph-level classification. *Mean*, *sum* and *global attention* aggregation schemes were tested, with *mean* aggregation attaining the best performance.

The readout network is a *MLP* that maps learned graph representation to activity prediction, and is trained jointly with the graph encoder. Batch Normalization and *ReLU* activation was used both in the graph encoder and in the readout network. Dropout was applied after each graph layer and readout layer. Linear warmup with cosine annealing was used as a learning rate scheduler.

#### Adversarial data augmentation

GNEprop leverages the FLAG framework^26^, that relies on “free” adversarial training^68^ and multi-scale augmentations. In this strategy, the input *x* is augmented with *M* adversarial perturbations, *δ*_*t*_ where *t* = 1, … *M*, such that the final optimization step is:

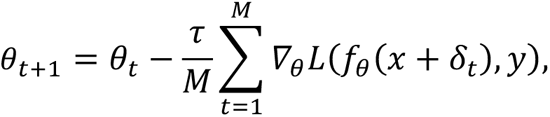

where *τ* is the learning rate, *L* is the binary cross entropy loss, *x* is a training sample and *y* is its label. We set *M* = 5. Perturbations *δ*_1:*M*_ are computed as follows: *δ*_1_ = *U*(−*α*, *α*), *δ*_*i*+1_ = *δ*_*i*_ + *α* ⋅ *sign*(*∇*_*θ*_*δ*_*i*_). The ascent step size is set *α* = 10^−3^. Therefore, augmentations have a maximum scale of *m* ⋅ *α*.

#### Self-supervised representation learning

During training, *N* graphs (minibatch) are input denoted by *G*^*k*^, with *k* = 1, …, *N*. Each graph was stochastically augmented twice, generating two augmented graphs 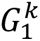 and 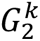. Structural augmentations^32,34^ (node dropping, edge dropping, edge perturbations, subgraph removal) and molecular augmentations^69^ were experimented with, attaining best results with subgraph removal (drop ratio = 0.25) in preliminary evaluations, which was used for following experiments. The model encoder *f*(⋅) and a non-linear projection head^70^ *g*(⋅) were jointly trained to distinguish whether a pair of augmented graphs come from the same graph or not, thus learning how to “undo” the augmentation pipeline. Negative samples were not explicitly sampled, but generated from other molecules in the same batch. The contrastive loss between 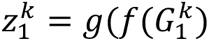 and 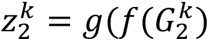 is defined through the normalized temperature-scaled cross entropy loss (NT-Xent)^25^:

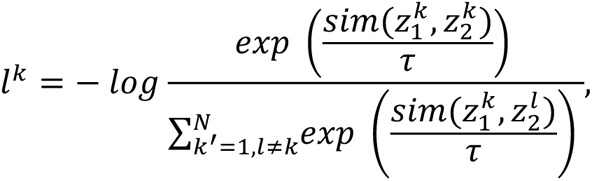

Where *τ* denotes the temperature parameter (here, *τ* = 0.1), and *sim* measures the similarity between embedded augmentations (here, *sim* is the cosine similarity). The projection head is a 3-layer *MLP*, with hidden dimension 512 and output dimension 256.

For pretraining, 122 million unique molecules from ZINC15^33^ previously identified in Stokes et al.^14^ as suitable antibiotic-like space were used. The dataset was split 90/5/5, resulting in 110M unique molecules for training. Validation metrics were monitored throughout training, and test molecules were used to visualize the learned representations (see below).

#### Meta-learning finetuning

After supervised finetuning starting from the self-supervised representations, the model is further finetuned through a meta-learning strategy to simulate domain shifts during training. This strategy is inspired by similar approaches that have been successfully applied in other domains, such as computer vision and reinforcement learning^29,71^. These methods have been shown to improve the robustness of the trained models to distribution shifts by exposing them to simulated shifts during training.

In this phase, the validation set is further split into a *meta-test* and a new *validation* set, in a 50/50 ratio. The validation set is used as before (in particular, for early stopping), while the meta-test set is used during training as described in the following. Each training update involves two steps. In the first step, the gradients are computed with respect to the original training set, *∇*_*θ*_*L*(*f*_*θ*_(*data*_*train*_)), leading to updated parameters *θ*′ after optimization. In the second step, the updated model is “virtually” evaluated on the meta-test set, and the loss *L*(*f*_-E_(*data*_*meta* *test*_)) is optimized *with respect to the original parameters θ*. The final loss is the weighted average of this new loss and the original loss on the train set, where the weight is a hyperparameter set to 0.3. The finetuning is performed for 5 epochs (where each epoch is defined as a full pass of the meta-test set), using the same hyperparameters of the original model, except for a constant learning rate (instead of warmup and annealing). In practice, the optimization leverages the MAML procedure^72^, computing the second-order optimization so that the improvement on the training set also improves the meta-test set. This strategy improved the AP on the full HTS dataset in the scaffold-cluster setting compared to only supervised finetuning, but led to comparable AP for the released dataset (potentially due to the reduced diversity and size of the meta-test, which could hinder generalization), and it has therefore been excluded from the latter.

#### Analysis of self-supervised representations

1M holdout molecules (<1% of the “antibiotic-like” space) were visualized as encoded with the pre-trained GNEprop. Learned molecular representations included 3,000 features. UMAP plots were created using the umap-learn python library with default parameters and Euclidean distance metric to reduce dimensions from 3,000 to 2. For each chemical property plot, the “spectral” colormap was used in matplotlib. Chemical properties were computed using RDKit functions*: Chem.Descriptors.MolWt()*, *Chem.Descriptors.TPSA(), Chem.Descriptors.MolLogP()* and *Chem.Descriptors.NumRotatableBonds().* Medicinal chemistry filters were computed using RDKit functions for computing chemical properties, and then are defined as follows: **Lipinski** (rule-of-five): MW <= 500, logP <= 5, h_bond_donor <= 5, h_bond_acceptors <= 10; **REOS**: 200 <= MW <= 500, -5 <= logP <= 5, 0 <= h_bond_donor <= 5 and 0 <= h_bond_acceptors <= 10, -2 <= formal_charge <= 2, 0 <= rotatable_bonds <= 8 and 15 <= heavy_atoms <= 50; **drug-like**: MW < 400, num_of_rings > 0, rotatable_bonds < 5, h_bond_donor <= 5, h_bond_acceptors <= 10, logP < 5; **antibiotic-like**^73^: 250 <= mw <= 550, tpsa >= 80, logp <= 2.0, h_bond_donor >= 2, h_bond_acceptors >= 4, rotatable_bonds >= 2. UMAP plots for Morgan fingerprints were created following Stokes et al.^14^: RDKit was used to compute a Morgan fingerprint for each molecule with radius = 2, number of bits = 2,048, and Jaccard (*i.e.*, Tanimoto) distance metric.

A linear evaluation protocol was followed in line with recent studies^74,75^. For the downstream evaluation task, three datasets from the MoleculeNet benchmark^36^ were used, choosing datasets in the physical chemistry category, as they are challenging properties to be predicted by structure-based methods and may be related to antibacterial activity: Lipophilicity (log*D*; 4,200 compounds), ESOL (water solubility; 1,128 compounds), and FreeSolv (experimental hydration free energies; 642 compounds). Each dataset was randomly split 30-fold 80/20. Test root-mean-squared-error (RMSE) was reported as mean and SEM over the folds. A Ridge regressor (*i.e.*, linear least squares with *L*2 regularization) was used as implemented in scikit-learn for the linear model. Ordinary least squares linear regression did not converge using Morgan fingerprints as input. All parameters of the Ridge regressor were kept as default, except for *α*, that was grid-searched and optimized for each experiment/model (tested values: [0.1, 1, 10, 100]). For GNEprop experiments, projection layers representations were combined, as they not only are more compact, but also proved to improve representation quality^70^. In particular, the input representation was *p*_1_ + *p*_<_, where *p*_3_ is the representation after the *i* projection layer. The resulting representation had 512 dimensions. For Morgan fingerprints experiments, RDKit was used to compute the fingerprint for each molecule.

#### Active learning pipeline

We investigated whether GNEprop can be used to support the acquisition of new training data in an active learning (AL) setting. Given that the size and diversity of the training data can have a significant impact on the model’s performance and the fact that new experiments are costly and time-consuming, an iterative strategy where the model is used to drive the selection of new compounds to be added to the training data can support virtual screening efforts. In the context of this study, these experiments help validate GNEprop predictions (both in terms of probability of activity and uncertainty estimation). We designed a simulated AL set-up, in which we started by randomly selecting a subset of the original training dataset and automatically expanding it over multiple acquisition rounds. The acquisition of new compounds is driven by the uncertainty of the model, estimated using several uncertainty estimation techniques, namely Gaussian processes^76^, Monte Carlo Dropout^77^, and Deep Ensembles^78^. For reference, we also considered a random selection baseline. Uncertainty estimation and active learning methods have been successfully applied in the context of deep learning-driven molecular property prediction in recent years^42,79,80^.

We considered different scenarios: *large* initial budget (in which we started from 5% of the training set randomly selected), *small* initial budget (in which we started from 0.1% of the training set randomly selected), and *all positive* initial budget (in which we started from all the active molecules, in addition to 0.1% of the negatives randomly selected). In addition to reporting the performance of the model at each iteration, we also report the fraction of positive samples recovered. In all experiments, we conducted 30 AL iterations (batched acquisition). Performance is averaged across 6 scaffold-based splits, and models have been trained for 30 epochs. For Monte Carlo Dropout, we used a dropout probability of 0.2 and 100 random samples. For Gaussian processes, we used a single SVDKL layer^81^ with hidden size of 256 instead of the original prediction layer, Gaussian prior *σ* = 0.1, and 100 samples used to estimate the uncertainty.

### Results on public antibacterial dataset

The dataset of 2,335 compounds from Stokes et al.^14^ was used for initial model training, optimization, and validation. Hyperparameter search was performed using Bayesian optimization, as implemented in the library Ax (https://ax.dev/), with 50 iterations of optimization, yielding: neural network hidden size = 500, number of message-passing steps = 5, number of readout layers = 1, dropout probability = 0.13, learning rate = 4.94e-05. All models were tested using scaffold splitting (as implemented in Yang et al.^37^) with 20 folds. Splits were conserved across all the experiments on this dataset to provide direct comparisons.

Several configurations of GNEprop were benchmarked (**Fig. 2c**). When a pre-trained encoder was not used, the encoder was randomly initialized with He initialization^82^. When RDKit features were passed, 200 additional molecule-level features, computed with RDKit, were passed to the learned molecular representation by concatenating them before the readout layers, following Stokes et al^14^. When using pretrained weights, a semi-supervised strategy was pursued: pretraining via self-supervised learning was followed by supervised fine-tuning. For the fine-tuning step, no parameter is frozen in the encoder, except for atom/bond embeddings and batch normalization.

For the comparison with chemprop, the same model and hyperparameters described in Stokes et al.^14^ were used. Parameters not specified were kept to their default values in the released chemprop software (http://chemprop.csail.mit.edu/). Chemprop was tested with additional molecular features computed by RDKit (as done in Stokes et al.^14^), and without them. Unlike in Stokes et al.^14^, results were obtained using a more challenging scaffold splitting strategy (as implemented in Yang et al.^37^), instead of a random splitting. For chemprop, hyper-optimization was re-run for the scaffold splitting setting to find the best hyperparameters that lead to similar performance. All evaluation metrics, except BEDROC, were computed using the scikit-learn library. In particular, AUPRC was calculated using the *average_precision_score* function. BEDROC was calculated using RDKit.

### Results on phenotypic HTS dataset

#### Scaffold-cluster splitting strategy

Scaffold-cluster splitting aims to simulate the real-world scenario where test molecules do not only correspond to novel scaffolds or variations, but entirely novel chemical spaces. All scaffolds were encoded using the pretrained encoder, then clustered following a community detection approach and pipeline^39^. Briefly, a *k*-nearest neighbors (*k*-NN) graph (*k* = 15) was computed between pairs of compounds using a Euclidean distance metric. Clusters of scaffolds were computed on the *k*-NN graph using the Leiden algorithm^83^. Each cluster was interpreted as a distinct molecular space, and used as a basic unit for splitting the dataset (*i.e.*, all the molecules with a certain scaffold were kept in the same split, such that the splitting strategy is still scaffold-based). UMAP embedding and Leiden clustering were performed with the implementation in scanpy^84^. All parameters were kept as default, except the Leiden resolution which controls the coarseness of the clustering, with higher values leading to more clusters. To balance between having enough clusters (for subsequent multi-folds splitting) and keeping meaningful, large groups, a resolution = 0.5 that led to 80 clusters was chosen, and allowed robust splitting. Clusters were randomly partitioned between training/validation/test in 70/15/15 proportions. Distributions for different splitting strategies were obtained computing the maximum Tanimoto similarity of Morgan fingerprints of all the training molecules for each molecule in the test set, for random, scaffold and scaffold-cluster splitting.

#### Models training

Models for HTS data were initially trained with the same hyper-parameters that yielded best performance on the public data, without further optimization. For the RF model, the same strategy as in Stokes et al.^14^ was followed: RDKit was used to compute the Morgan fingerprint for each molecule, with radius = 2 and number of bits = 2,048. Additionally, Morgan fingerprints with radius up to 3 and number of bits up to 4,096 were tested, without observing statistically significant improvements over the initial configuration.

#### Activity cliffs analysis

There were 498791 unique scaffolds^85^ in the HTS dataset, 181,435 of which include more than one molecule, and 2045 include at least one antibacterial and one non-antibacterial molecule. The difference in activity within these 2045 scaffold families was defined as activity cliffs. In total, the 1,255 scaffold families with activity cliffs included 217424 molecules (median mols/scaffold = 12, mean mols/scaffold = 106.3 molecules, antibacterial/scaffold ≈ 1.85). Analysis was performed on families with activity cliffs using the same models without further optimization. Importance scores were computed using the Integrated Gradients (IG)^41^ method, following a strategy similar to Sanchez-McCloskey et al.^86^: the input dataset was augmented with null graphs (*i.e.*, molecules with the same topology but with zeroed node and edge features) of 30% of positively-labeled samples, which had a 50% probability of having a positive or negative label. Doing this, a baseline molecular graph *̄* was generated from a graph *x*, by zeroing atom/bond features, and with an expected output ∼0.5. As both atom and bond features were used as input to the GNN, the attribution score was computed for each atom and bond feature, and atom/bond attributions were then obtained by aggregating over atoms/bonds. To explain antibacterial compounds, negative attributions were ignored after aggregation (*i.e.*, only features that are deemed important to classify a molecule as antibacterial were highlighted). Finally, atom and bond features were linearly rescaled in the [0,1] range, and importance scores were visualized using a viridis colormap.

#### Cell membrane permeability

The compounds antibacterial against the *E. coli* mutant strain were assayed against a wild type *E. coli* strain, to identify molecules which can also penetrate the outer membrane. Of the 3,090 initial hits, only 3,074 were available for the second screening round: 2,768 compounds were antibacterial only against the mutant strain (label: "inhibiting") and 306 molecules were antibacterial against both mutant and wild type (label: "cell-penetrating"). Compounds with < 25 μm IC50 were labeled as WT antibacterial. GNEprop was retrained using the same procedure described for the main screening, but in a three-class setting (“inantibacterial”, “inhibiting”, “cell-penetrating”). In this case, as cell-penetrating molecules are rare, it was beneficial to use a class-weighted cross-entropy loss to account for imbalanced data. To learn differential features between inhibiting and cell-penetrating compounds, the model was re-trained in a two-class setting. Scaffold splitting, was used 80/10/10 for train/validation/test sets, for 20 folds. Notably, unsupervised pre-training was critical to differentiate between the two conditions (without pretraining: AUROC = 0.715 ± 0.015, AUPRC = 0.304 ± 0.017; with pretraining: AUROC = 0.745 ± 0.014, AUPRC = 0.335 ± 0.019). To explain cell-penetrating molecules, the same IG-based pipeline used to explain activity cliffs was applied.

### Virtual screening inference

GNEprop has been applied to the “REAL Database” provided by Enamine (2020 version), including 1.4 billion compounds. The REAL (REadily AccessibLe) space includes enumerated synthetically accessible compounds. The same model described for the retrospective results has been re-trained with a 95/5/0 scaffold-cluster splitting (i.e., omitting the test set to maximize data usage). Eight replicates have been trained, resampling the validation set in each in an ensembling strategy. All replicates have been used to screen the REAL Database, averaging the predicted probabilities. Scaling GNEprop across 64 A100 GPUs, we were able to screen the whole space for all replicates in less than 48 hours. In the end, compounds with a predicted probability greater than 0.6 were selected, which constitute the virtual hits (44,437 molecules). For all virtual hits, the maximum similarity to training actives and known antibiotics has been computed. Tanimotor similarity on Morgan fingerprints (radius = 2, number of bits = 2,048) as implemented in RDKit has been used.

This smaller set of virtual hits is manageable through standard computational tools, and it has been further refined to identify the final set. In particular, compounds with unwanted functional groups (n=1,100) and higher similarity to known antibiotics (n=1,092) have been discarded. The remaining lightly filtered compounds have been clustered to promote diversity and explore multiple hypotheses within the fixed budget. Sphere Exclusion clustering with Atom-Atom-Path similarity (AAP)^87^ (AAP radius = 0.25) was used, resulting in 984 clusters. Clusters with very low predicted kinetic solubility (cKinSol) (i.e., <2.0 µM) have been manually analyzed. When all compounds in the clusters have low cKinSol (n=52), they were discarded. Otherwise, clusters were kept selecting the compound with the highest cKinSol. In all other cases, the compound with the highest predicted activity by GNEprop has been selected for each cluster. This led to 932 selected molecules. From this set, 503 molecules have been manually prioritized for synthesis. 345 molecules have been successfully synthesized.

### Novel MOA detection

#### Antibiotic clustering

A dataset of 196 known antibiotics was assembled, grouped into 31 classes according to their target. These antibiotics were embedded through the learned self-supervised encoder; clusters of antibiotics were defined by using the annotated targets. The resulting clusters were quantitatively compared to those obtained using conventional Tanimoto similarity on Morgan fingerprints through different clustering metrics (Silhouette^88^, Calinski-Harabasz^89^, Davies-Bouldin^90^ and Dunn index^91^). GNEprop representations led to better clustering according to all the considered metrics.

#### Novelty detection

To identify whether a compound belongs to a *known* or a *novel* mechanism of action, the problem was formulated as an Out-Of-Distribution (OOD) detection task, where a model needs to distinguish between novel samples coming from known or novel classes^50^. In this case, each molecule is a sample and targets correspond to classes. In computer vision, representations learned through contrastive learning can boost OOD detection performance. Thus, the methodology proposed in Winkens et al.^51^ was extended to the graph contrastive framework. In particular, the OOD problem was framed as a density estimation problem: for each known class *c*, an *n*-dimensional multivariate Gaussian *N*(*μ*_*c*_, *Σ*_*c*_) was estimated based on known samples; a *novelty score s* was then computed for each unseen sample, with respect to the class-conditional Gaussian distributions, based on the Mahalanobis distance:

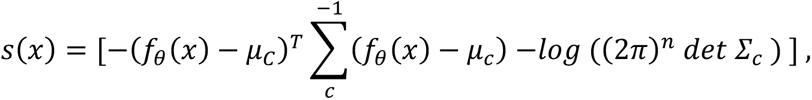

where *f*_-_is the encoder and (*μ*_*c*_, *Σ*_*c*_) denote, respectively, the mean and covariance matrix of the multi-variate Gaussian distribution for class *c*. A low *s*(*x*) indicates that an unseen compound lies far from all known classes in the embedding space, and is thus more likely to represent an OOD example.

In addition to Winkens et al.^51^, and similarly to Lee et al.^50^, sample representations were obtained by ensembling representations (unweighted sum) across multiple projection layers to improve their expressiveness. To validate the proposed approach, all classes were randomly split in two sets: known (*K*) and unknown (*U*) classes. The test set included all samples from the *U* classes and an equal number of samples randomly selected from the *K* classes. The training set included all the remaining samples from the *K* classes. Through this setting, the OOD problem can be seen as a balanced binary classification problem, and evaluated through standard classification metrics (AUROC, AUPRC) on the test set. The training and test sets were randomly re-sampled 20 times to report robust statistics. Additionally, the number of holdout classes *U* was varied from 1 to 3, to assess the performance across different settings. This approach was benchmarked against three baselines, each of them following a similar strategy, but where one or more components were replaced with alternatives: (1) replacing the Mahalanobis distance from class distributions with Euclidean distance from the centroids of each cluster (Euclidean-centroids baseline); (2) using Morgan fingerprints instead of GNEprop representations (MF-OOD baseline); or (3) using Morgan fingerprints and Tanimoto distance from class centroids. All the baselines were evaluated through the exact same splits and metrics used to evaluate the proposed methodology. Results indicate that both the learned representations, and using Mahalanobis distance from class distributions, are critical to best performance.

## Acknowledgements

We acknowledge Yiming Xu, Christopher Heise, Maureen Beresini, Timothy Dawes, Jennifer Perry-Ruzic, Jessica Hankins, Mary Kate Alexander, Jingyu Diao, Nicholas Nickerson for designing, executing, and enabling the phenotypic screen.

## Author contributions

STR, MWT, AR, GS and TB designed the study. GS and TB designed GNEprop, with the help of ZL, KC, ND, JCH, JML, MK, and YB. STR and AM conducted the HTS and collected the data. GZ and ND optimized and scaled up the GNEprop model. GS, STR, ZL, NS, KRB, and TB designed and conducted the virtual screen. KRB and STR conducted the experimental validation and biological characterization of compounds. NS, KRB, ES, ZL, and GS analyzed the data. All authors wrote the manuscript and provided feedback.

## Competing Interests

GS, STR, ZL, KRB, NS, KC, ND, JCH, JF, LG, ES, AR, MWT and TB are employees of Genentech, Inc. and shareholders of Roche. YB is advisor for Recursion Pharmaceuticals.

---

**Extended Data Figure 1.**
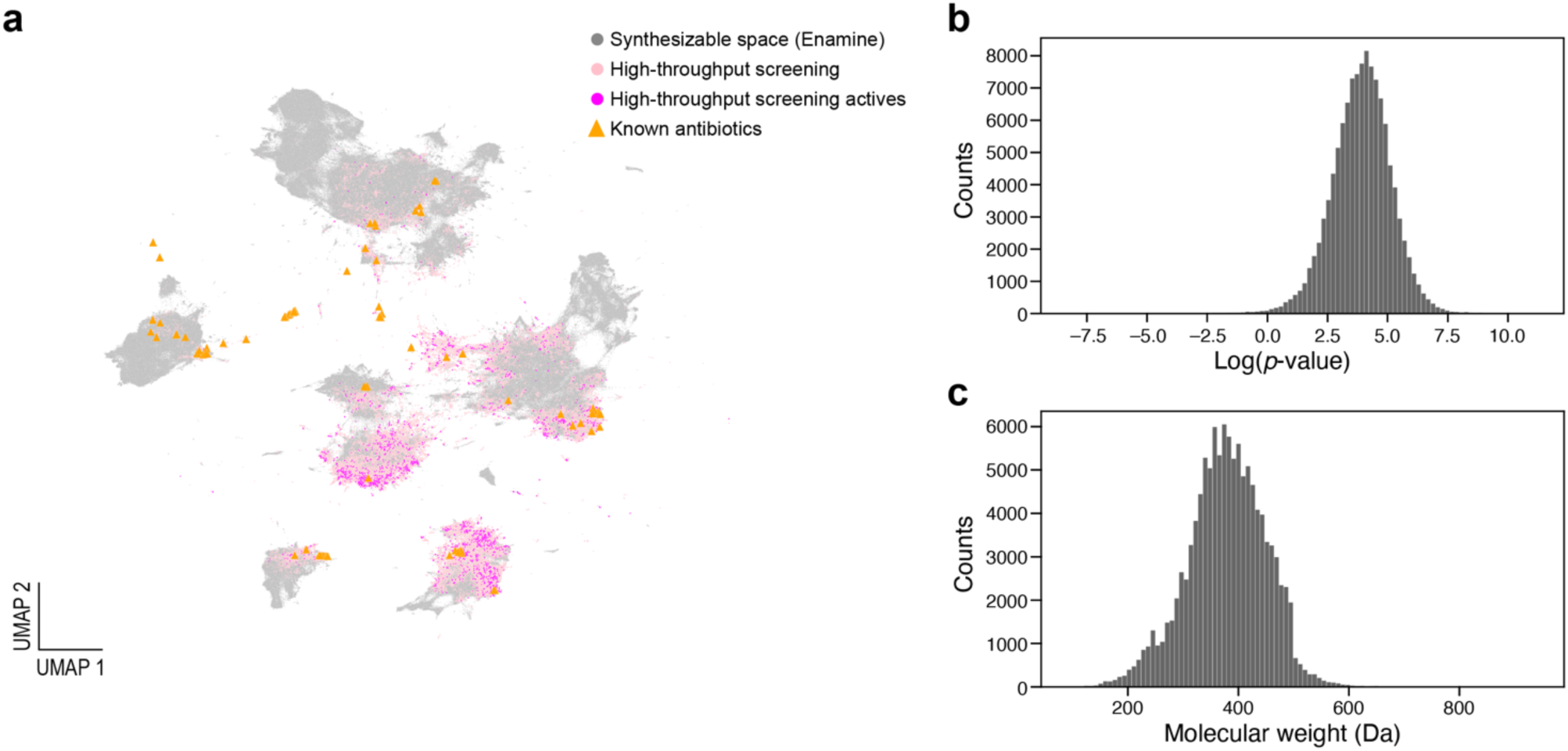
The newly-released *GNEtolC* dataset includes over 115,000 compounds, encompassing ∼3,000 antibacterial compounds. **a**, UMAP. Visualization of the dataset, compared to (randomly sampled) Enamine space and known antibiotics. Each dot is a molecule colored by the source (legend, **Methods**). Representations are based on self-supervised GNEprop representations and Euclidean distance. **b**, distribution of logP for molecules in the *GNEtolC* dataset. **c**, distribution of molecular weight for molecules in the *GNEtolC* dataset.

**Extended Data Figure 2.**
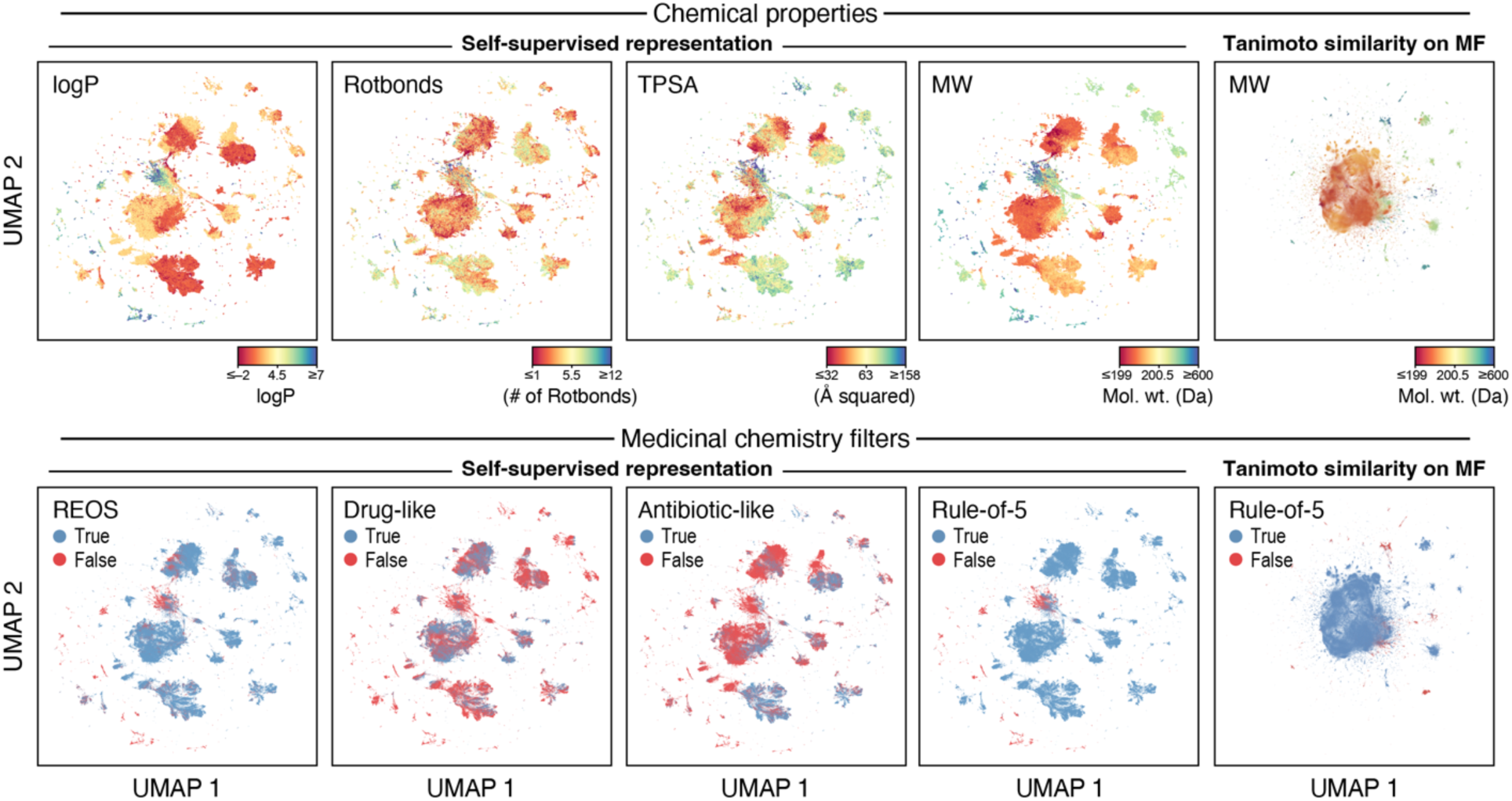
Self-supervised representations for additional properties and medicinal chemistry filters of compounds.

**Extended Data Figure 3.**
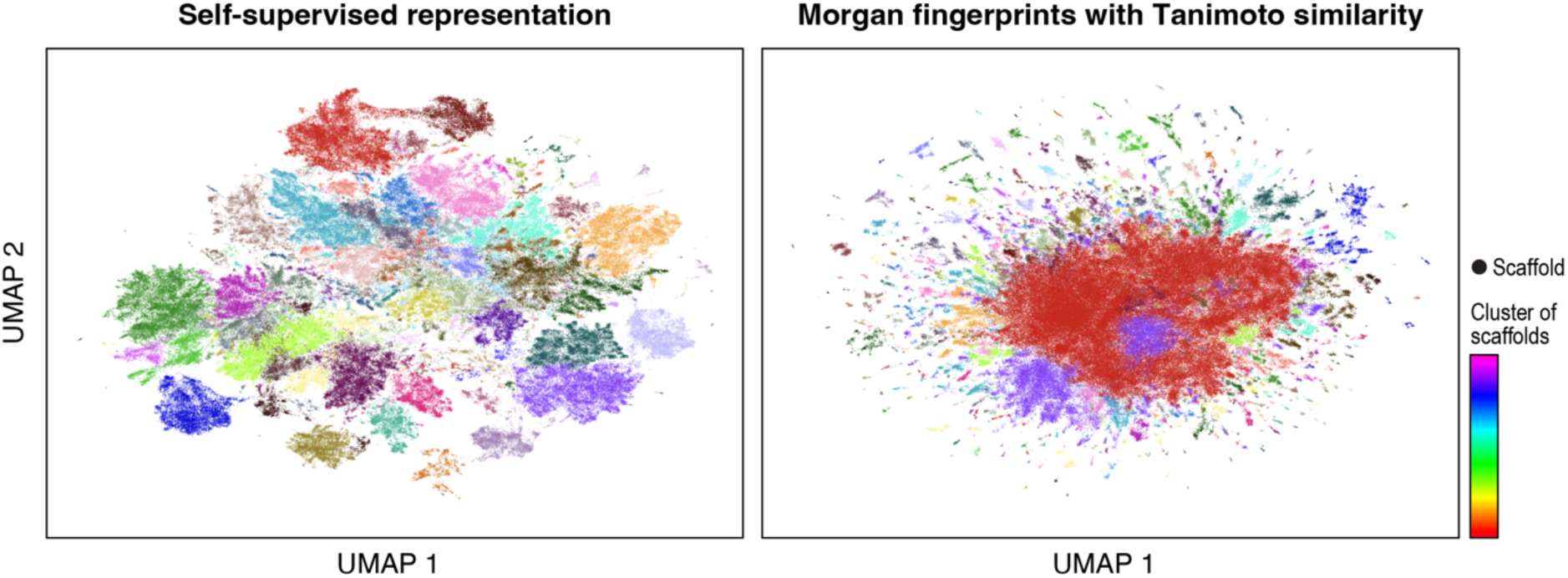
Comparison of clusters derived from learned representations and Morgan fingerprints for molecules in the HTS dataset. Scaffolds are encoded through self-supervised representations (left) and Morgan fingerprints with Tanimoto similarity (right). A *k*-nearest neighbor (*k*-NN) graph followed by the Louvain community detection algorithm is used to derive clusters (colors). Self-supervised representations lead to meaningful clusters that can be used as a unit for splitting the dataset. In contrast, Morgan fingerprints do not lead representations that can be easily clustered.

**Extended Data Figure 4.**
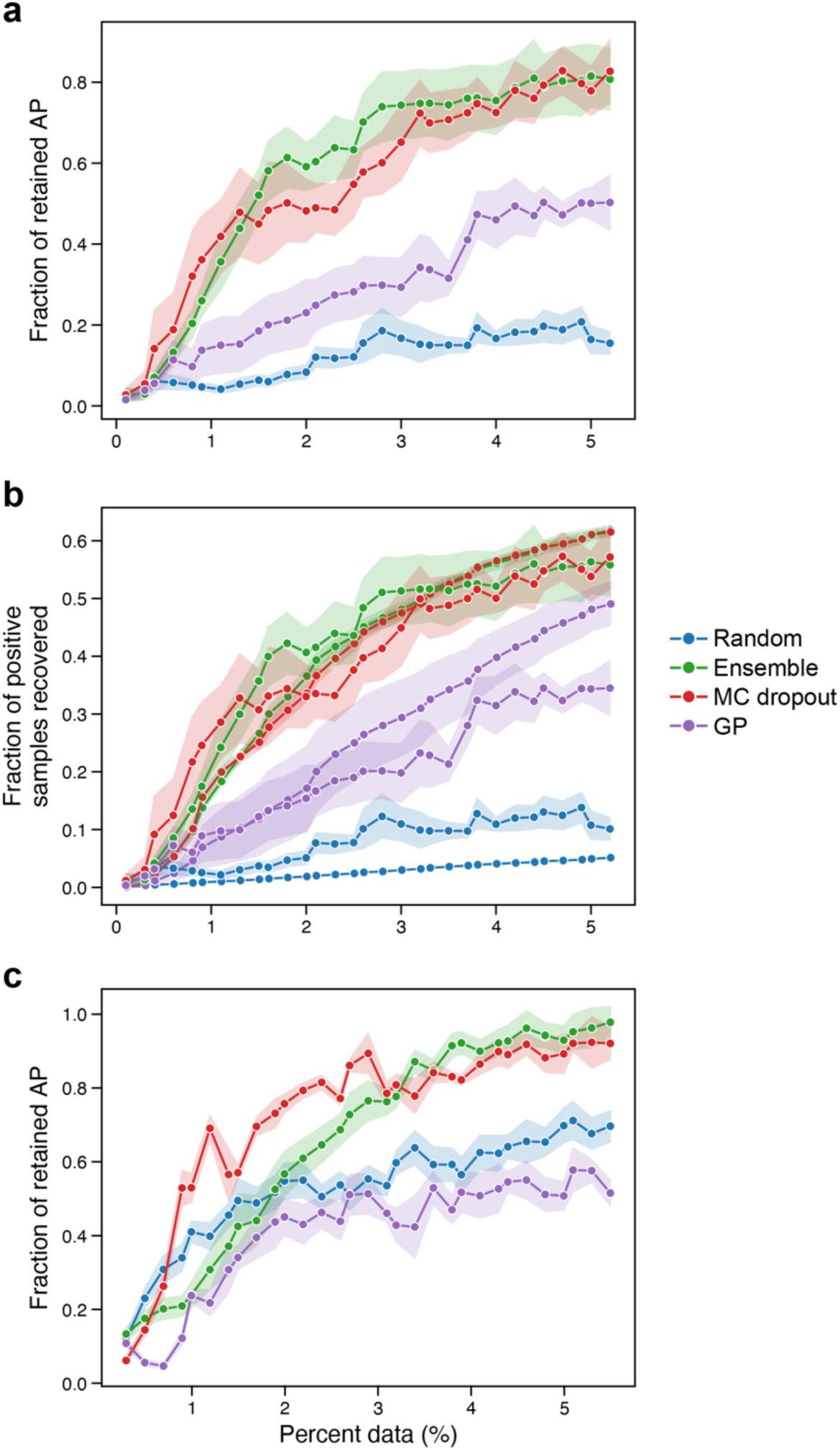
Additional active learning results. **a**, the fraction of retained AP and **b**, the fraction of positive samples recovered as a function of the percentage of the data used in the active learning setting. **c**, the fraction of retained AP in the setting where all positives are added to the initial training set. In all panels, colors denote different uncertainty estimation methods, dots denote experiments and shade area denotes standard error.

**Extended Data Figure 5.**
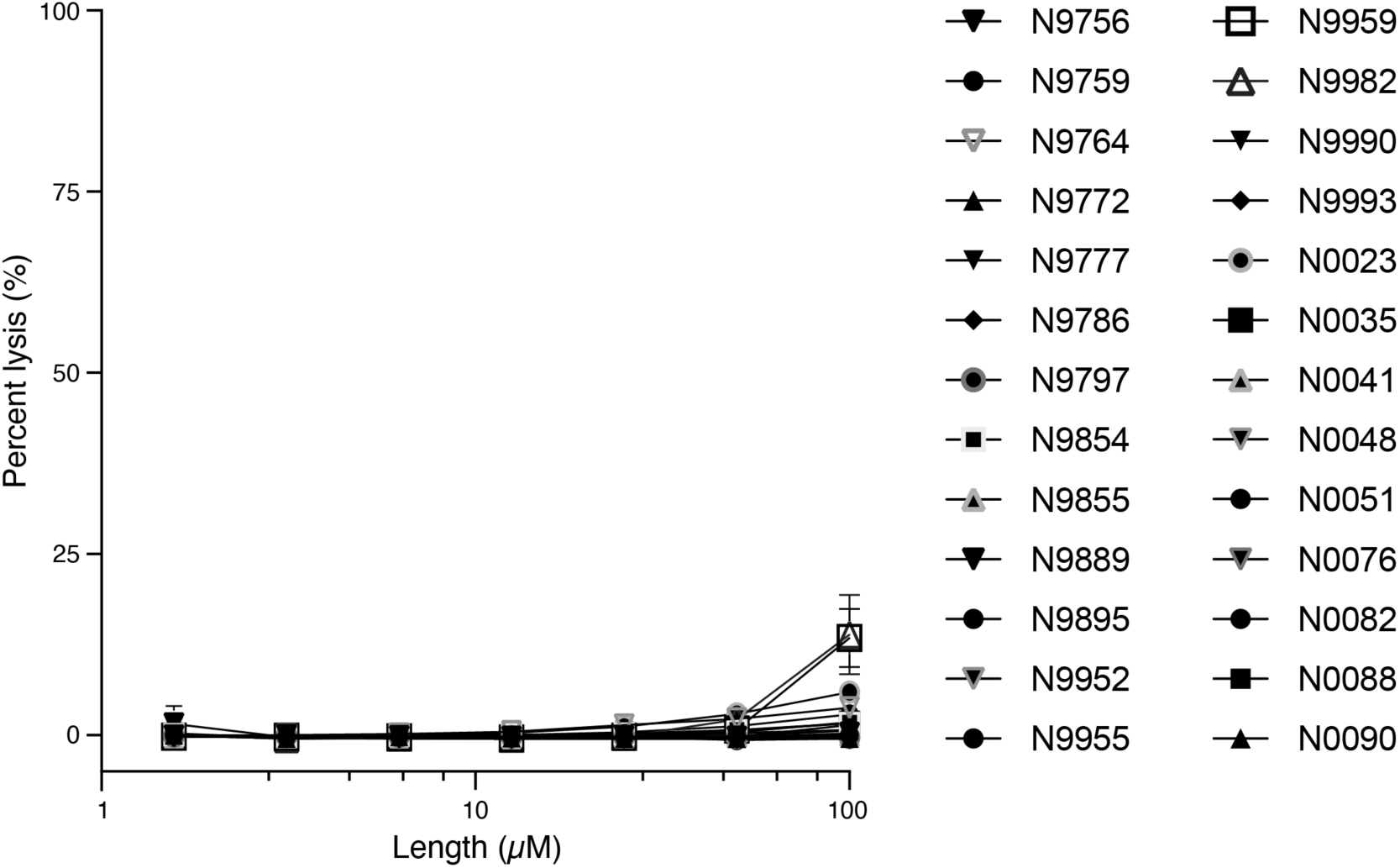
Red blood cell lysis assay. Compounds at indicated concentrations were evaluated for hemolytic activity after 18 hours. Percent lysis of samples calculated relative to zero and 100% lysis samples (methods). Shown are data from 3 biological replicates, mean +/- SD.

**Extended Data Figure 6.**
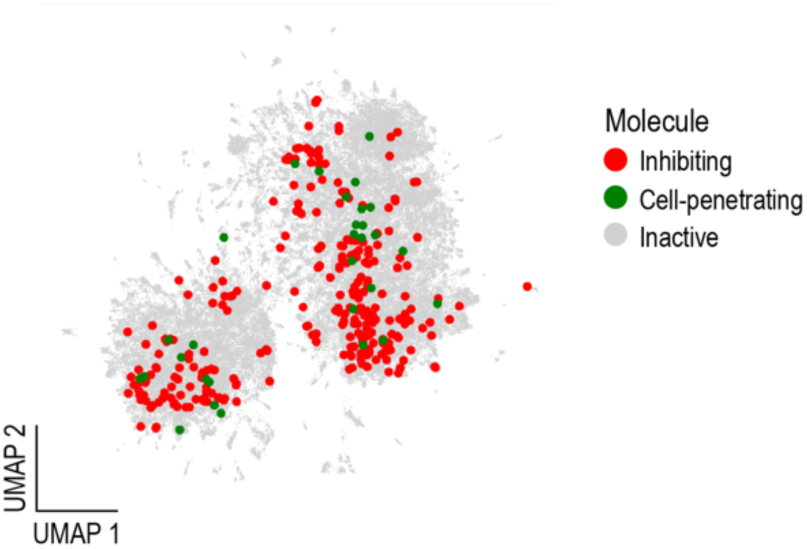
Low dimensional plot of self-supervised representation showing inactives (grey dots), actives on *E. coli* Δ*tolC* (red dots) and actives on wild type *E. coli* (green dots).

**Extended Data Table 1. GNETolC dataset.**

See file GNEtolC.csv

**Extended Data Table 2.** Results on public dataset.

| Model | AUROC | AUPRC | BEDROC<sub>$\alpha=20$</sub> | Precision | Recall | F1 |
| --- | --- | --- | --- | --- | --- | --- |
| **Train/test via scaffold splitting** |  |  |  |  |  |  |
| chemprop (no RDKit features) | 0.821 ± 0.018 | 0.429 ± 0.041 | 0.534 ± 0.040 | 0.505 ± 0.074 | 0.276 ± 0.049 | 0.318 ± 0.047 |
| chemprop (+ RDKit features) | 0.838 ± 0.018 | 0.447 ± 0.037 | 0.565 ± 0.032 | 0.578 ± 0.062 | 0.317 ± 0.042 | 0.377 ± 0.045 |
| GNeprop (no pretraining; no RDKit features) | 0.822 ± 0.019 | 0.491 ± 0.031 | 0.617 ± 0.024 | 0.658 ± 0.051 | 0.346 ± 0.038 | 0.439 ± 0.042 |
| GNeprop (no pretraining; + RDKit features) | 0.846 ± 0.014 | 0.514 ± 0.032 | 0.633 ± 0.026 | 0.641 ± 0.043 | 0.365 ± 0.036 | 0.445 ± 0.036 |
| GNeprop (+ pretraining; no RDKit features) | **0.859 ± 0.016** | **0.570 ± 0.030** | 0.671 ± 0.028 | 0.687 ± 0.037 | **0.437 ± 0.018** | **0.513 ± 0.034** |
| GNeprop (+ pretraining; + RDKit features) | 0.852 ± 0.018 | 0.565 ± 0.034 | **0.679 ± 0.029** | **0.749 ± 0.038** | 0.414 ± 0.039 | **0.512 ± 0.037** |
| **Train/test via random splitting** |  |  |  |  |  |  |
| chemprop (no RDKit features) | 0.848 ± 0.020 | 0.502 ± 0.041 | 0.604 ± 0.040 | 0.453 ± 0.064 | 0.374 ± 0.053 | 0.395 ± 0.052 |
| chemprop (+ RDKit features) | 0.875 ± 0.013 | 0.563 ± 0.026 | 0.660 ± 0.026 | 0.725 ± 0.049 | 0.421 ± 0.039 | 0.500 ± 0.03 |
| GNeprop (no pretraining; no RDKit features) | 0.893 ± 0.011 | 0.649 ± 0.030 | 0.738 ± 0.024 | **0.771 ± 0.041** | 0.479 ± 0.036 | 0.570 ± 0.029 |
| GNeprop (no pretraining; + RDKit features) | 0.898 ± 0.012 | 0.653 ± 0.028 | 0.742 ± 0.024 | 0.726 ± 0.033 | 0.511 ± 0.036 | 0.587 ± 0.029 |
| GNeprop (+ pretraining; no RDKit features) | **0.917 ± 0.010** | **0.671 ± 0.027** | **0.755 ± 0.021** | 0.715 ± 0.035 | 0.524 ± 0.037 | 0.590 ± 0.030 |
| GNeprop (+ pretraining; + RDKit features) | 0.915 ± 0.010 | 0.668 ± 0.029 | 0.750 ± 0.024 | 0.712 ± 0.036 | **0.536 ± 0.039** | **0.598 ± 0.033** |

**Extended Data Table 3.** Results on HTS dataset.

| Model | AUROC | AUPRC | BEDROC<sub>$\alpha=20$</sub> | Precision | Recall | F1 |
| --- | --- | --- | --- | --- | --- | --- |
| **Scaffold splitting** |  |  |  |  |  |  |
| Random forest | 0.853 ± 0.009 | 0.167 ± 0.023 | 0.531 ± 0.024 | 0.654 ± 0.077 | 0.033 ± 0.008 | 0.062 ± 0.014 |
| GNeprop | **0.927 ± 0.004** | **0.302 ± 0.025** | **0.651 ± 0.016** | **0.694 ± 0.029** | **0.204 ± 0.022** | **0.311 ± 0.029** |
| **Scaffold-cluster splitting** |  |  |  |  |  |  |
| Random forest | 0.763 ± 0.009 | 0.060 ± 0.013 | 0.357 ± 0.020 | **0.472 ± 0.119** | 0.010 ± 0.005 | 0.019 ± 0.009 |
| GNeprop | **0.877 ± 0.007** | **0.126 ± 0.019** | **0.488 ± 0.014** | **0.467 ± 0.053** | **0.077 ± 0.015** | **0.130 ± 0.024** |

**Extended Data Table 4.** Results on GNEtolC dataset (GNEprop model).

|  | Scaffold splitting | Scaffold-cluster splitting |
| --- | --- | --- |
| AUROC | 0.728 ± 0.01 | 0.651 ± 0.024 |
| AUPRC | 0.244 ± 0.022 | 0.146 ± 0.025 |
| BEDROC<sub>$\alpha=20$</sub> | 0.387 ± 0.02 | 0.267 ± 0.03 |
| Precision | 0.504 ± 0.053 | 0.463 ± 0.062 |
| Recall | 0.192 ± 0.019 | 0.097 ± 0.021 |
| F1 | 0.27 ± 0.02 | 0.155 ± 0.03 |

**Extended Data Table 5.** Results on HTS dataset, activity cliffs only.

| Model | AUROC | AUPRC | BEDROC<sub>$\alpha=20$</sub> | Precision | Recall | F1 |
| --- | --- | --- | --- | --- | --- | --- |
| Scaffold splitting |  |  |  |  |  |  |
| Random forest | 0.798 ± 0.013 | 0.232 ± 0.033 | 0.448 ± 0.025 | 0.625 ± 0.160 | 0.018 ± 0.005 | 0.036 ± 0.010 |
| GNEprop | **0.885 ± 0.010** | **0.354 ± 0.033** | **0.557 ± 0.021** | **0.723 ± 0.036** | **0.162 ± 0.027** | **0.258 ± 0.038** |
| Scaffold-cluster splitting |  |  |  |  |  |  |
| Random forest | 0.719 ± 0.014 | 0.129 ± 0.019 | 0.312 ± 0.023 | 0.579 ± 0.153 | 0.006 ± 0.003 | 0.012 ± 0.005 |
| GNEprop | **0.797 ± 0.016** | **0.215 ± 0.022** | **0.404 ± 0.022** | **0.583 ± 0.091** | **0.052 ± 0.014** | **0.094 ± 0.024** |

**Extended Data Table 6. Full list of confirmed hits.** 44,437 compounds are selected by the GNEprop model (label: “Virtual hit”), among which 345 compounds are further selected for in-house experiment (label: “Selected for in-house experiment”), and finally 82 are tested active in single-dose experiments (label: “Confirmed hit”).

See file screening_hits.xlsx

**Extended Data Table 7.** Dose-response assay against *Saccharomyces cerevisiae*. Table shows compounds with <30 µM IC50.

| Compound | *S. cerevisiae* | *E. coli* WT | *E. coli* ΔtolC | *S. aureus* |
| --- | --- | --- | --- | --- |
| N9757 | 3.9 | 6 | 0.56 | 1 |
| N9761 | 7.6 | 40 | 4 | 3.1 |
| N0046 | 17.0 | 40 | 7.1 | 2.6 |
| N0027 | 17.0 | 6.6 | 2 | 2 |
| N9831 | 18.0 | 26 | 1.4 | 1.6 |
| N9939 | 22.0 | 40 | 3.4 | 1.9 |

**Extended Data Table 8.** Minimal inhibitory concentration assay against *E. coli ΔtolC*, *E. coli,* and *Staphylococcus aureus*.

| Compound | *E. coli* ΔtolC | *E. coli* WT | *S. aureus* |
| --- | --- | --- | --- |
| N9756 | 40 | >40 | >40 |
| N9759 | 5 | >40 | 10 |
| N9764 | >40 | >40 | >40 |
| N9772 | 20 | >40 | >40 |
| N9777 | 0.313 | >40 | >40 |
| N9786 | 10 | >40 | >40 |
| N9797 | 20 | >40 | >40 |
| N9854 | 2.5 | >40 | 10 |
| N9855 | >40 | >40 | >40 |
| N9889 | 20 | >40 | 40 |
| N9895 | 10 | >40 | >40 |
| N9952 | 0.625 | >40 | >40 |
| N9955 | 1.25 | >40 | 2.5 |
| N9959 | 2.5 | >40 | 5 |
| N9982 | 1.25 | >40 | 2.5 |
| N9990 | 5 | >40 | 20 |
| N9993 | >40 | >40 | >40 |
| N0023 | 2.5 | >40 | 2.5 |
| N0035 | 40 | >40 | 40 |
| N0041 | 5 | >40 | 5 |
| N0048 | 10 | >40 | 40 |
| N0051 | 5 | >40 | 1.25 |
| N0076 | >40 | >40 | >40 |
| N0082 | 1.25 | >40 | 1.25 |
| N0088 | 20 | >40 | 40 |
| N0090 | 2.5 | >40 | 20 |

**Extended Data Table 9.** N9777 and N9786 resistant E. coli K-12 MG1655 strains. Only variants present in at least 50% of mapped reads at the site of variation are reported, and the frequency of each variant is indicated in parentheses. Synonymous substitutions are not reported.

| Compound | Strain | Non-synonymous substitutions and indels |
| --- | --- | --- |
| N9777 | Mutant 1 | LpxH (b0524): F141L (100%) |
| N9777 | Mutant 2 | LpxH (b0524): F128L (100%), UbiG (b2235): D13N (100%) |
| N9786 | Mutant 1 | FabZ (b0180): P59T (100%) |
| N9786 | Mutant 2 | FabZ (b0180): R121S (100%) |
| N9786 | Mutant 3 | FabZ (b0180): F21L (100%) |
| N9786 | Mutant 4 | FabZ (b0180): P104L (100%), YieL (b3719): D269E (100%) |
| N9786 | Mutant 5 | FabZ (b0180): T2_T3insTN (100%) |
| N9786 | Mutant 6 | FabZ (b0180): T2_T3insTN (100%) |
| N9786 | Mutant 7 | AcpP (b1094): D57E (100%) |
| N9786 | Mutant 8 | FabZ (b0180): E48D (100%) |

**Extended Data Table 10. Dataset of known antibiotics and targets.**

See file Extended_data_table_antibiotics.csv

**Extended Data Table 11.** Clustering scores for known antibiotics and targets.

| Model | Silhouette ↑ | Calinski-Harabasz ↑ | Davies-Bouldin ↓ | Dunn index ↑ |
| --- | --- | --- | --- | --- |
| GNeprop representations | **0.23** | **21.97** | **1.57** | **0.21** |
| Morgan fingerprints | 0.11 | 16.60 | 1.70 | 0.09 |

**Extended Data Table 12.** Novel target detection results.

| Model | AUROC | AUPRC |
| --- | --- | --- |
| 1 excluded target |  |  |
| Euclidean-centroids | 0.73 ± 0.08 | 0.84 ± 0.04 |
| MF-OOD | 0.85 ± 0.06 | 0.92 ± 0.03 |
| Tanimoto-centroids | 0.80 ± 0.08 | 0.87 ± 0.05 |
| GNeprop-OOD | **0.97 ± 0.01** | **0.98 ± 0.01** |
| 2 excluded targets |  |  |
| Euclidean-centroids | 0.88 ± 0.03 | 0.90 ± 0.02 |
| MF-OOD | 0.84 ± 0.04 | 0.90 ± 0.03 |
| Tanimoto-centroids | 0.83 ± 0.04 | 0.86 ± 0.04 |
| GNeprop-OOD | **0.91 ± 0.02** | **0.94 ± 0.01** |
| 3 excluded targets |  |  |
| Euclidean-centroids | 0.76 ± 0.04 | 0.81 ± 0.03 |
| MF-OOD | 0.85 ± 0.02 | 0.89 ± 0.02 |
| Tanimoto-centroids | 0.84 ± 0.04 | 0.84 ± 0.04 |
| GNeprop-OOD | **0.90 ± 0.02** | **0.92 ± 0.02** |

**Extended Data Table 13.** Strains and plasmid used in this study.

| Strain | Resistance gene | Source | GNEID |
| --- | --- | --- | --- |
| *E. coli* MG1655 | n/a | In-house | 2 |
| *E. coli* MG1655ΔtolC | Kan-R | In-house | 31 |
| *S. aureus* USA300 | n/a | ATCC (BAA-1556) | 23 |

| Plasmid | Description | Resistance gene | Source |
| --- | --- | --- | --- |
| pRP1195 | luxBADCE operon | Carb-R | * |

\*Plaut, R. D., Mocca, C. P., Prabhakara, R., Merkel, T. J. & Stibitz, S. Stably luminescent *Staphylococcus aureus* clinical strains for use in bioluminescent imaging. PLoS One 8, e59232 (2013).

